# Detection and classification of long terminal repeat sequences in plant LTR-retrotransposons and their analysis using explainable machine learning

**DOI:** 10.1101/2024.06.11.598549

**Authors:** Jakub Horvath, Pavel Jedlicka, Marie Kratka, Zdenek Kubat, Eduard Kejnovsky, Matej Lexa

**Affiliations:** Faculty of Informatics, Masaryk University, Botanicka 68a, 60200 Brno, Czech Republic; Department of Plant Developmental Genetics, Institute of Biophysics of the Czech Academy of Sciences, Kralovopolska 135, 61200 Brno, Czech Republic; National Centre for Biomolecular Research, Faculty of Science, Masaryk University, Kamenice 5, 625 00 Brno, Czech Republic

**Keywords:** eukaryote, repeat, transposable elements, deep learning, CNN-LSTM, DNABERT, sequence analysis, regulatory mechanisms, transcription factor binding sites, TFBS

## Abstract

**Background:** Long terminal repeats (LTRs) represent important parts of LTR retrotransposons and retroviruses found in high copy numbers in a majority of eukaryotic genomes. LTRs contain regulatory sequences essential for the life cycle of the retrotransposon. Previous experimental and sequence studies have provided only limited information about LTR structure and composition, mostly from model systems. To enhance our understanding of these key compounds, we focused on the contrasts between LTRs of various retrotransposon families and other genomic regions. Furthermore, this approach can be utilized for the classification and prediction of LTRs.

**Results:** We used machine learning methods suitable for DNA sequence classification and applied them to a large dataset of plant LTR retrotransposon sequences. We trained three machine learning models using (i) traditional model ensembles (Gradient Boosting - GBC), (ii) hybrid CNN-LSTM models, and (iii) a pre-trained transformer-based model (DNABERT) using k-mer sequence representation. All three approaches were successful in classifying and isolating LTRs in this data, as well as providing valuable insights into LTR sequence composition. The best classification (expressed as F1 score) achieved for LTR detection was 0.85 using the CNN-LSTM hybrid network model. The most accurate classification task was superfamily classification (F1=0.89) while the least accurate was family classification (F1=0.74). The trained models were subjected to explainability analysis. SHAP positional analysis identified a mixture of interesting features, many of which had a preferred absolute position within the LTR and/or were biologically relevant, such as a centrally positioned TATA-box, and TG..CA patterns around both LTR edges.

**Conclusions:** Our results show that the models used here recognized biologically relevant motifs, such as core promoter elements in the LTR detection task, and a development and stress-related subclass of transcription factor binding sites in the family classification task. Explainability analysis also highlighted the importance of 5’- and 3’-edges in LTR identity and revealed need to analyze more than just dinucleotides at these ends. Our work shows the applicability of machine learning models to regulatory sequence analysis and classification, and demonstrates the important role of the identified motifs in LTR detection.

## 1 Background

Long terminal repeats (LTRs) are essential regulatory sequences of retrotransposons and retroviruses, often found in high copy numbers in many eukaryotic genomes (Baucom et al. 2009, Klaver and Berkhout 1994). LTR retrotransposons are the main repeat type in most plant genomes (Jedlicka et al. 2020, Luo et al. 2022). While retrotransposons propagate through transcription and subsequent insertion, experimental methods for studying transposable elements are limited due to the inactivation of a majority of the genomic copies in most of the life cycle except for reproductive cells and in response to stress (Bennetzen and Wang 2014, Grandbastien et al. 2005, Sigman and RK. 2016). In addition, experiments are typically only carried out on a small number of model sequences and organisms.

Genomic sequence analysis can thus provide important additional information about the composition, classification, and function of LTRs in LTR retrotransposons and even in their evolutionarily contrasting element subtypes (superfamilies and families, see Wicker et al. (2007)). This approach has shown some success when applied to full-length LTR retrotransposon sequences in plants (Arango-López et al. 2017), including machine learning approaches (Orozco-Arias et al. 2022), but has not been applied specifically to LTRs whose structure is more loosely defined than the structure of internal coding regions of the retrotransposons. This imprecise characterization complicates the analysis of plant LTR sequences with traditional methods.

In a way, LTRs are “the closest cousins” of regulatory sequences such as promoters and enhancers. First, LTRs themselves function as promoters in transcription of their own LTR retrotransposon copy (Casacuberta and Santiago 2003), not unlike what happens in human LTR retroviruses, such as HIV (Dutilleul et al. 2020). They can drive the transcription of neighboring genes (Cui and Cao 2014). Second, there is ample evolutionary evidence that LTR-TEs contribute to the makeup of older regulatory sequences either by inserting into them, nearby, or providing the initial building material for subsequent regulation (Thompson et al. 2016). Both LTRs and gene regulatory sequences (promoters, enhancers), have an increased ability to bind transcription factors (Hermant and Torres-Padilla 2021).

In LTR retrotransposons, it is relatively easy to delineate the LTRs since they occur in two copies, one at each end of the transposable element (TE), and in the case of bona-fide insertions also carry tandem site duplications (TSDs) at their outer boundaries (Turcotte et al. 2001). However, their internal composition is often difficult to unravel. Functional LTRs must always contain three regions important for the life cycle of the entire TE. These are known as U3, R and U5, and can be determined experimentally (Arkhipova et al. 1986). U3 is known to bind regulatory proteins important for transcription and its components are capable of serving both as enhancers and promoters. U5 may contain additional regulatory signals and it borders on or partially overlaps the primer binding site (Zhang et al. 2014). The R region is delineated by the transcription start and termination sites. Region identification by in-silico sequence analysis is problematic. Sequences of plant LTRs are variable not only in sequence composition but also in their length, ranging from around a hundred bps to several thousands (Du et al. 2010). We are looking for ways in which sequence analysis can shed light on to the internal structure of LTRs and identify regulatory regions, such as transcription factor binding sites (TFBS) and their type and absence/presence in different TE families.

Deep learning (Sapoval et al. 2022) and transformer-based models (Vaswani et al. 2017) have the potential to address these challenges, having been successfully applied in recent genomic data analyses (Ji et al. 2021, Jumper et al. 2021), including the classification of full length LTR retrotransposons (Chen et al. 2024). While this approach demonstrated high classification accuracy, the learning process reflecting the biological features of LTR retrotransposon sequences has not yet been fully examined. Here we have employed these models for LTR sequence identification and classification, focusing on model interpretability as a tool to extract both existing and new biological knowledge about these regulatory sequences.

Due to the highly variable length and sequence composition of LTR sequences, LTR identification using common bioinformatics solutions poses a complicated problem. Machine learning (ML) methods can provide insight into complex relationships within the data with minimal prior assumptions due to the process of learning on input features. This allows us to uncover previously unrecognized properties, from the successful interpretation of the learned internal structure of such models. Here, we will focus on the state-of-the-art ML and deep learning (DL) mehodologies that have already proven useful in similar scenarios.

The Gradient Boosting classifier (GBC) is an ensemble-based method which iteratively trains multiple weaker learners on the pseudo-residuals of learners from previous iterations, with the goal of improving upon their prediction errors. The accuracy of the model is dependent on its hyperparameters, such as the number of sub-estimators used, as well as the specific hyperparameters of the sub-estimators. This can be improved by techniques that identify an optimal combination of these parameters for a given dataset. In general, the ensemble model tends to be relatively robust to overfitting and achieves good results in fairly complex biological tasks (Kotov et al. 2023, Messad et al. 2019).

The combination of convolutional neural networks (CNN) and LSTM nodes has proven efficient both in natural language processing tasks and in the biological domain (Gunasekaran et al. 2021, Liang et al. 2020). The effectiveness of this combination stems from the ability of convolutional filters to capture local patterns, including, but not limited to, those of TFBS and the ability of the LSTM to recognize remote dependencies and the co-existence of these patterns. LSTM nodes are able to selectively filter out information about the input sequence through the use of a gating mechanism. This enables the LSTM network to retain relevant information, discard irrelevant details, and carry over crucial context from previous elements in the sequence (Hochreiter and Schmidhuber 1997).

The BERT family of models (Devlin et al. 2018) is a relatively recent tool that has seen many successful applications, mainly in natural language processing but also recently in the challenge of transposable element classification (Chen et al. 2024). The BERT model is a transformer-based neural network model that utilizes the mechanism of attention to recognize the context of words and embed sentences into a fixed-size vector. Such embeddings have a number of key properties, which make them useful for further downstream tasks. One example is that semantically similar sentences tend to have embeddings whose cosine distance is small. An important feature of popular BERT-based models is their pre-trained nature, meaning that fine-tuning to custom data requires much smaller datasets, making it also much faster than training from scratch. One such model pre-trained on DNA sequences is DNABERT. Analogical to natural language, the function of DNA is also based on its internal structure and the order of its sub-features, making the DNABERT model an attractive candidate when dealing with variable length sequences with unknown structure.

Machine learning and deep learning have seen many advancements in the area of model interpretability techniques, promoting better comprehension beyond the black-box approach that particularly deep learning models have been known for. Appreciation of how a model makes decisions will give us a better understanding of our data and the ability to detect class-specific features. Such applications provide a way to pinpoint key structural properties of data such as DNA sequences, where the order of elementary features defines a certain biological function. The techniques used in this work range from the direct analysis of the model structure such as the analysis of convolutional layer filters, to more complex algorithmic tools such as SHAP (Lundberg and Lee 2017).

As machine learning and deep learning have already been successfully used to delineate promoters and TF binding sites An et al. (2022) and lncRNAs (Danilevicz et al. 2023) in genomic sequences, we set out to investigate here their ability to improve our understanding of LTR structure, modularity and genomic sequence composition. We also wanted to know whether models based on different algorithmic principles would show any shortcomings or advantages in regulatory sequence analysis.

## 2 Results

As mentioned above, our aim was to specifically analyze plant LTR sequences that are more variable and dynamic, and therefore more difficult to study than coding regions of LTR retrotransposons. Due to their inherent modularity as promoters and higher variability in overall length, comparing and clustering them via sequence alignment is more complicated. An alternative approach appears to be motif identification. Dedicated software, such as MEME (Bailey and Elkan 1994) has often been successful in extracting motifs from promoters. However, the absence of expression data makes the location of common motifs much more difficult.

Therefore, we tested a simple model using k-mers of annotated LTRs (data from Zhou et al. (2021) further described and used throughout the paper), where we compared k-mers (k = 6) using the Jaccard Similarity Index (JSI) between the TE super-families - Ty1/Copia and Ty3/Gypsy (Supplementary Figure 1A). These superfamilies share 37 % of unique k-mers. Furthermore, the dendrogram of individual LTRs created from their JSI values did not reliably distinguish representatives of the two superfamilies (Supplementary Figure 1B). Additionally, we compared the occurrence of k-mers in LTRs with the length of corresponding non-LTR sequences from the genomes of relevant plants as controls (Supplementary Figure 1C). The results indicate that in more than 80 % of plant species (56 out of 69), the JSI value of k-mers present in LTR and non-LTR sequences was higher than 75 %. Clearly, neither sequence alignment, nor simple k-mer counting are powerful enough to meaningfully cluster LTR sequences from different families or species, or to isolate the subsequences responsible for their specific function.

In response to these shortcomings of the above-mentioned classical approaches, we decided to employ forms of machine learning that allow flexibility both in terms of the motifs being sought (we may not necessarily know them in advance) and their relative or absolute positions within the LTR.

The main focus of our work was to discover features of plant LTRs that contribute the most to the ability of a machine learning model to detect or classify LTR nucleotide sequence of plant LTR retrotransposons. Figure 1 shows the overall data flow and tools used here, which consisted of the input data collection and filtering, its preparation and encoding for subsequent use in ML models, model construction and learning, the final interpretation of the results, and identification of the most influential features.

**Figure 1.**
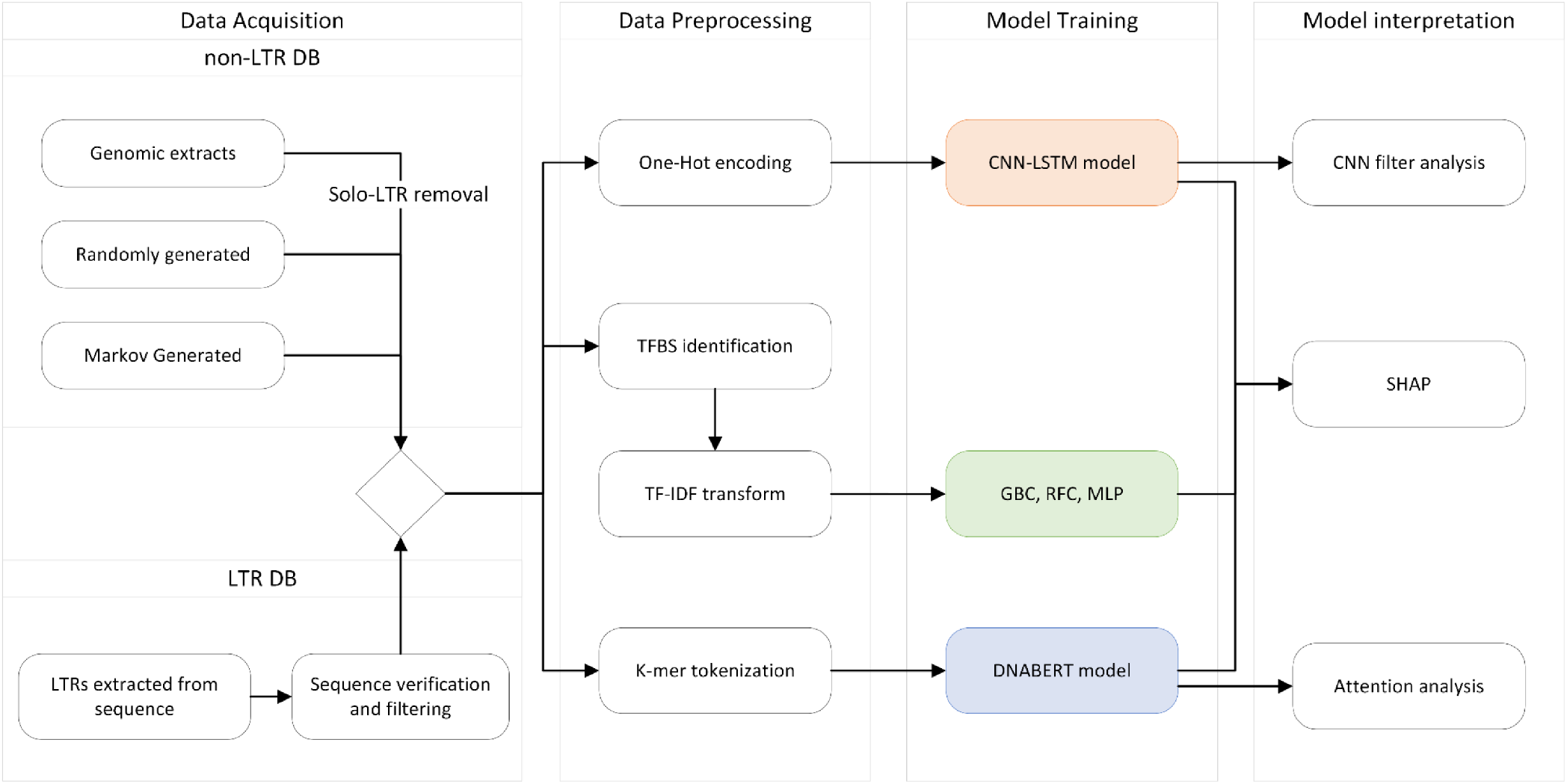
Data processing and computational workflow diagram. Input DNA sequences (positive and negative LTR sets) were pre-processed for the three alternative modeling approaches (to obtain TF binding site presence, one-hot encoding, and k-mers). The last two columns show the software tools used in individual branches of the analysis.

Three main tasks of this study were: (i) LTR detection - learning to distinguish LTR and LTR-negative sequences; (ii) Superfamily classification - learning to distinguish Ty3/Gypsy and Ty1/Copia LTRs; and (iii) Family classification - classification into any of the 15 families selected for the input data.

Three types of machine learning models were employed for the above DNA sequence classification tasks (see section 3.3 of Methods for more details). First, a conventional ML model that uses TFBS counts as input, specifically a Gradient Boost classifier (GBC). Second, a hybrid CNN-LSTM network trained on one-hot encoded sequences and finally, a pre-trained, transformer-based DNABERT model (Ji et al. 2021) further trained on k-mer-tokenized sequences.

Recently several large-scale studies of plant LTR retrotransposons have been carried out. To provide us with a sufficiently high number of complete annotations we chose a study by Zhou et al. (2021) that produced “a comprehensive annotation dataset of intact LTR retrotransposons of 300 plant genomes”. Altogether, 2,593,685 LTR retrotransposons are available in this dataset, however after applying additional criteria for quality and redundancy, we ended up with 176,917 LTR retrotransposons (and their corresponding LTR pairs) from 75 plant species (see section 3.1.3 in Methods). To facilitate machine learning, an LTR-negative sequence dataset was prepared as described in section 3.1.2 in Methods.

### 2.1 Model training

The three types of models were trained on the input data as described in section 3.4 of Methods. The accuracy of the trained models as evidenced by computed F1 characteristics was in the range 0.68-0.89 (Table 1). Binary classifications of LTRs and their superfamily membership were easier to learn, while the hybrid CNN-LSTM model was the best overall. It is evident that learning just on JASPAR TFBS (in GBC) leads to lower accuracy, especially in LTR detection (F1=0.73 v. 0.85; difference of 0.12), while the differences in family and superfamily classification tasks are much lower (difference of 0.06-0.07). The lower differences in accuracy between models at the family level could reflect the biological fact that all LTR retrotransposons share the common elements important for their life cycle (and also for their detection and classification at higher levels), while less information is available from features recruited by individual families. More detailed results of the learning step are available in Supplementary Figures 2-5.

**Table 1.**
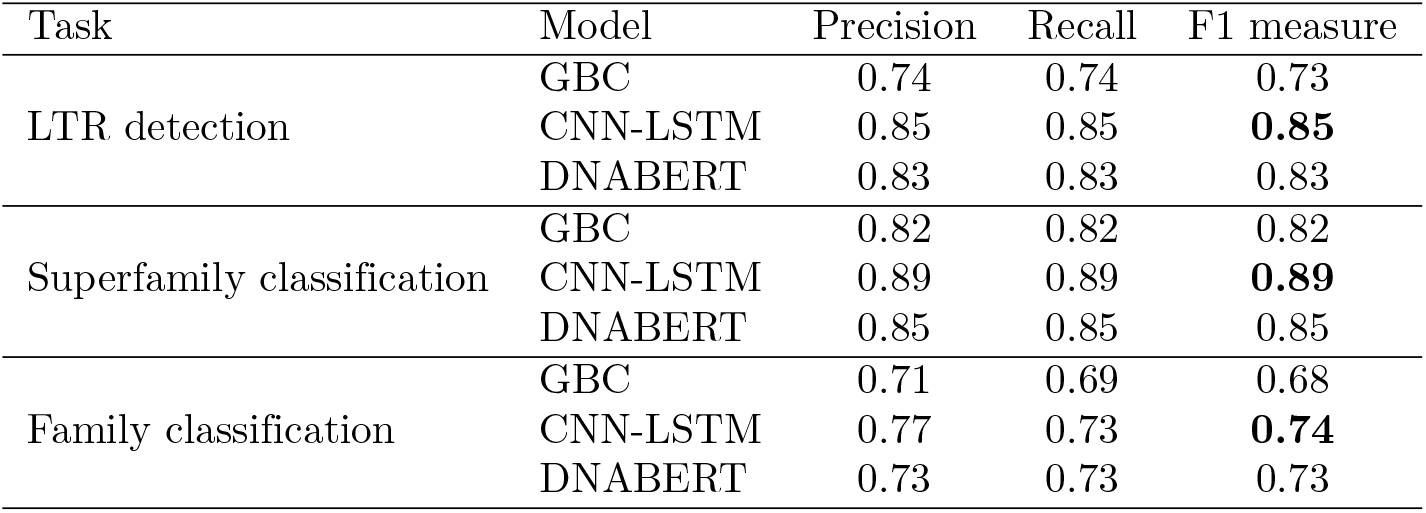
Precision, recall and F1 measure of three types of models in the three tasks. **GBS** - Gradient Boosting classifier; **CNN-LSTM** - a hybrid network model; **DNABERT** - pre-trained BERT. The best F1 values for a given task are shown in bold.

The superiority of the CNN+LSTM hybrid network model can be clearly seen in the family classification task (Figure 2). Despite having generally lower recall in the three most numerous families (Ale, CRM, Tekay), this network had however maintained higher precision than the other models (see also Supplementary Figures 6 and 7).

**Figure 2.**
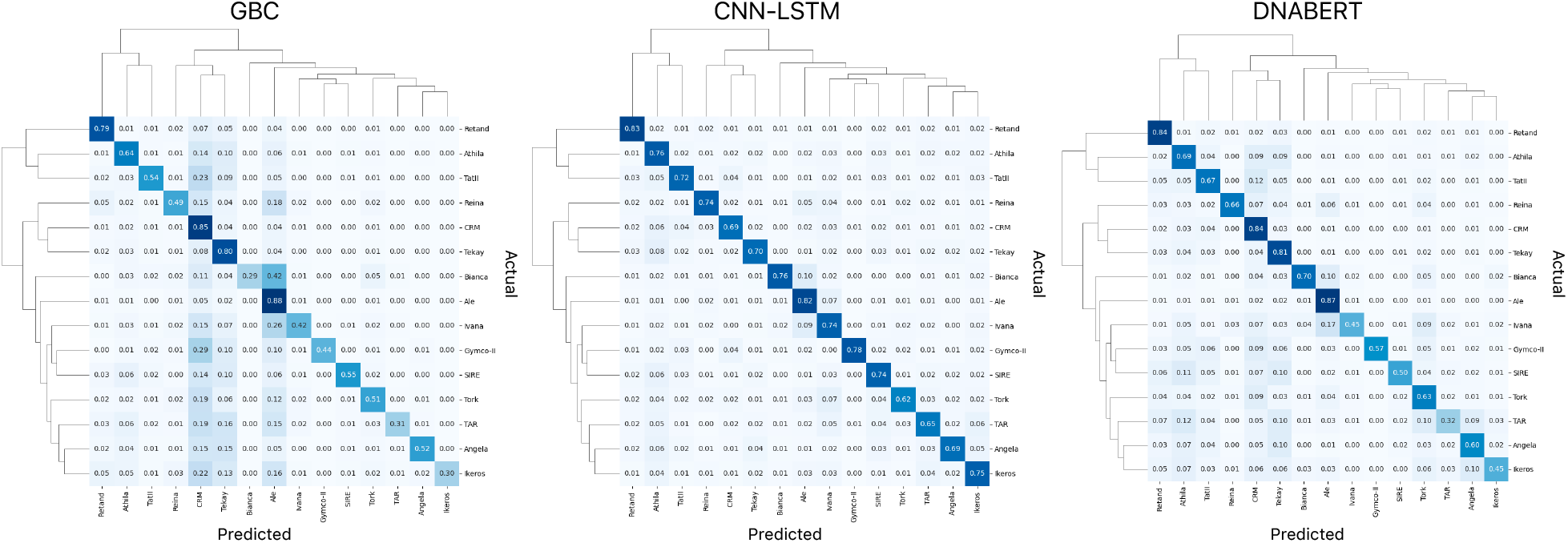
Cross-accuracy of the three model types in the family classification task. GBS - Gradient Boosting classifier; CNN-LSTM - a hybrid network model; DNABERT - pre-trained BERT.

### 2.2 Model interpretation

While the models trained to detect and classify LTRs can be useful in themselves, they largely represent black box models that provide little understanding of how these classifications actually materialized. This is a well-known and universal problem of machine-learning, and particularly deep-learning methods. Current deep learning approaches try to address this problem by specialized post-training analysis of the model and its inputs and outputs. We adapted two such approaches to the LTR classification problem presented here, some of which can only be used on specific model types. Derivation of Shapley additive explanations (SHAP) is by principle a model-agnostic method and can be applied to all models. Convolutional filter analysis was used for neural network models. The methods for interpretable machine learning used here are described in more detail in the Methods section 3.5.

#### 2.2.1 LTR

SHAP explanations were used in two modes on the GBC model to gauge the effect of TFBS in the analyzed DNA sequences. Jaspar TFBS were also used to visualize the trained filters in CNN models. The filters from the model were compared to Jaspar matrices using TomTom (Gupta et al. 2007). Finally, the DeepExplainer and Explainer modules of the SHAP package (Lundberg and Lee 2017) were used to calculate SHAP values along the analyzed LTR sequences in CNN and DNABERT models respectively, with the ambition of uncovering regions of LTRs most contributing to successful learning.

##### SHAP TreeExplainer (GBC)

First we analyzed models trained to recognize LTRs (as opposed to other random, or genomic sequences)(Figure 3,4). SHAP analysis of the GBC model shows the top 20 most impactful transcription factor binding sites (TFBS) present in LTR sequences and contributing most towards their classification (Figure 3a). To get an indication about which TFBS, or what kind of regulation might be specific to LTR retrotransposons in plants, we ran this set through gProfiler GOSt functional analysis (Kolberg et al. 2023). Apart from many hits to general TF-related terms (such as nucleus, DNA-binding, transcriptional regulation, etc.), the biological process results have also shown interesting subsets, namely: (i) response to stimulus, (ii) anatomical and multicellular development, and (iii) reproductive process (Supplementary Figure 8-10).

**Figure 3.**
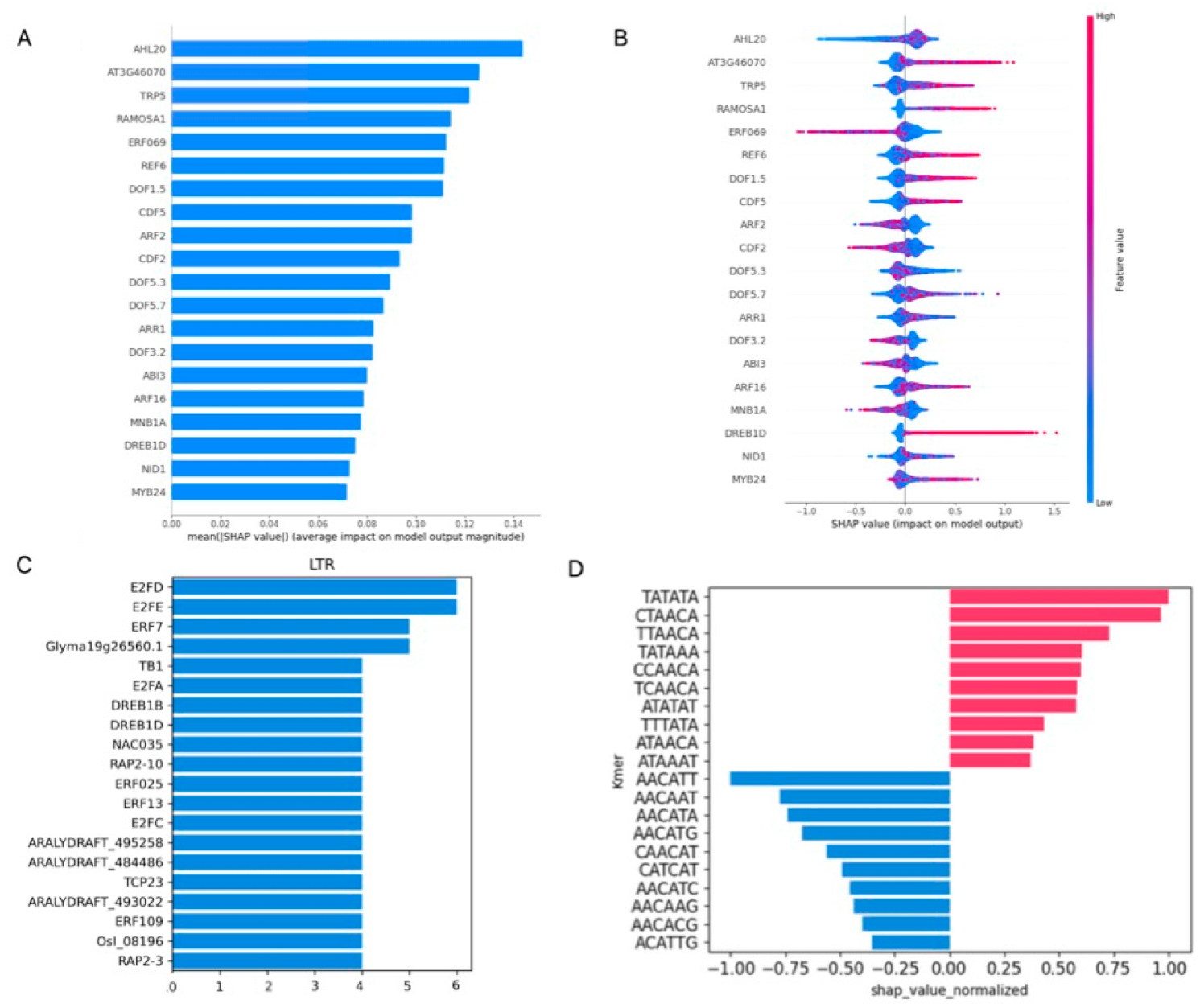
Main results of explainability analysis carried out on trained LTR detection models. **A** - Top twenty transcription factor binding sites (TFBS) with the highest mean SHAP value contribution (as input features) to the GBC model performance in LTR classification. Accross all LTRs the presence or absence of these TFBS was most important for classification of sequences as LTRs by the model (compared to other TFBS). **B** - Beeswarm plot showing the extent to which input features influenced model output. Color codes for the TF-IDF transformed occurrences of that particular TFBS as described in Methods section 3.2.1, horizontal axis shows SHAP values, indicating whether the effect of a TFBS presence in the analyzed sequence was positive, or negative. **C** - TomTom hits of first-layer CNN filters on JASPAR Core 2022 database. The X axis represents the number of CNN filters that were mapped to the specific TFBS as described in Methods, Section 3.5.2. **D** - Top contributions of individual k-mers to DNABERT classification as determined by SHAP analysis. Positively valued k-mers influence the classification towards the LTR class, while negative values influence the classification towards the non-LTR class.

**Figure 4.**
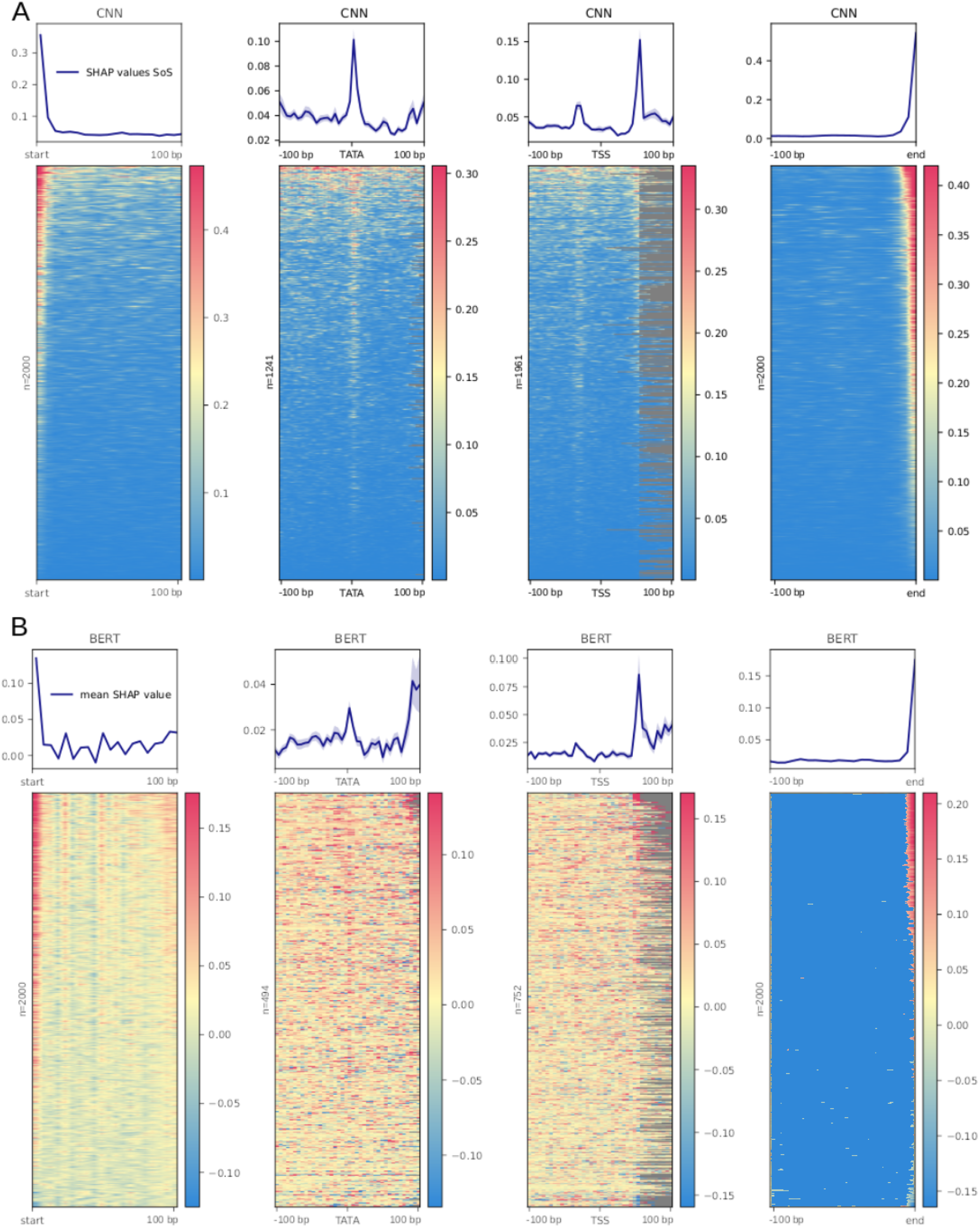
SHAP analysis of trained LTR detection models. k-mer based SHAP values were calculated along individual LTR sequences. To visualize their alignment between different sequences, sequences were aligned (from left to right) by their first base (start), predicted TATA box (TATA), predicted transcription start site (TSS), and their last base (end). Averaged SHAP values are shown as a line graph above, individual sequence values are color-coded. **A** - CNN model. **B** - DNABERT model

**Figure 5.**
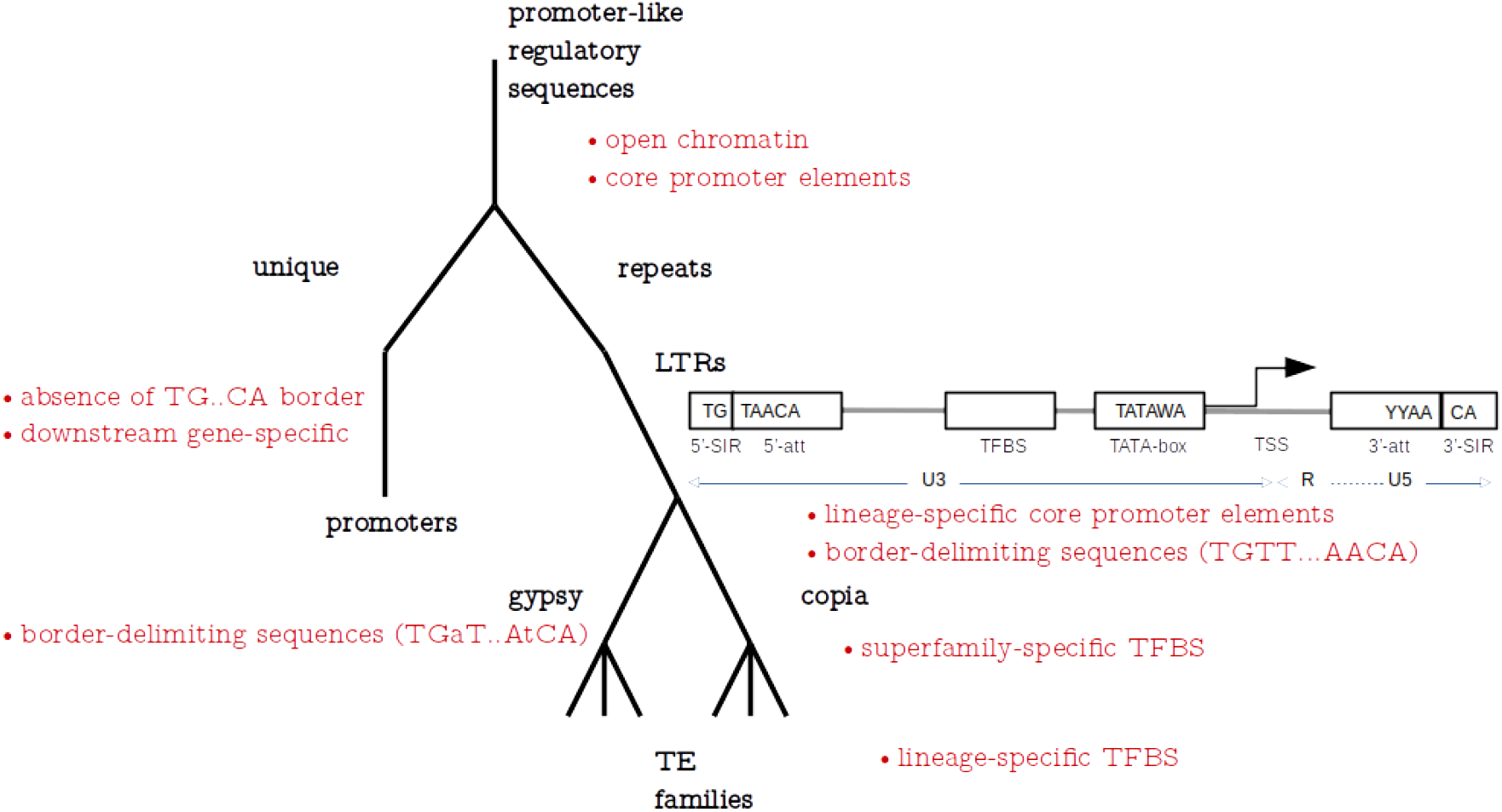
A hierarchy of promoter-like regulatory elements (including LTRs). **black** - subsets of regulatory sequences that ML models are trained on; **red** - sequence motifs specific for respective subsets that the models can use to classify the group correctly. Block diagram shows the structure of a typical LTR with sequence motifs assembled from k-mers discovered by the DNABERT model trained here

Beeswarm plots of the same analysis (Figure 3b) differentiate between the positive and negative contribution of the individual TFBS to classification. Specifically, ERF069 and DREB1D show a particularly sharp boundary of high/low SHAP values and positive/negative classification, although in opposite directions. ERF069 is an ethylene responsive element mostly absent from LTRs, while DREB1D on the other hand, is a dehydration responsive element that is associated with LTRs. A complete list of evaluated TFBS and their performance in the LTR classification task using the GBC model is provided as Supplementary File 1.

##### Filter analysis (CNN)

Another opportunity to look at TFBS as instrumental in LTR classification was via filter analysis of the CNN model. Figure 3c shows the top 20 results from the comparison of first-layer convolutional filters to JASPAR database TFBS, sorted based on the number of motif hits. gProfiler GOSt functional analysis shows results similar to the above paragraph, however in this case, only response to stimulus was present as a subset of TF-specific terms, embodied by the only common hit with the above analysis in DREBD1B. Filter analysis brought up several E2F and ERF family members’ binding sites. While the former are cell-cycle progression regulators, the latter are ethylene responsive factors that may have been used by the model as a negative indicator of LTR sequences (analogically to SHAP results in GBC above). A complete list of evaluated filters/TFBS and their presence as filters in the CNN model is provided as Supplementary File 2.

##### DeepExplainer (CNN)

The DeepExplainer module was implemented to visualize the location of sequence positions with the highest SHAP values in the CNN-LSTM model (Figure 4a). To visualize the alignment of possible signals between different sequences, the sequences were aligned by their first base (start), predicted TATA box, predicted transcription start site (TSS), and their last base (end) and shown as line graphs and heatmaps. In both the averaged line graph values and the heatmaps, three signals pop up as locations instrumental in LTR classification. They are the first few and the last few bases of the LTRs, as well as the TATA box predicted with TSSPlant Shahmuradov et al. (2017). It is also apparent in the CNN model, that the 5’ ends of the LTRs have a higher density of informative k-mers than their 3’ ends, possibly reflecting a typical TFBS position upstream of the TSS.

##### Explainer (k-mers, DNABERT)

Similarly to the SHAP analysis on TFBS and sequence positions, the analysis can be carried out on kmer-based models to identify k-mers present in LTR sequences that, in a given instance contribute more significantly to the classification of the sequence as an LTR or a non-LTR. The top 20 k-mers for the LTR classification task via the DNABERT model are shown in Figure 3d. We have identified overlaps among the k-mers that indicated their origin from a wider sequence motif (Supplementary File 3) and found the following putative consensus motifs to be present: (i) TATA[AT]A (positive SHAP values, a likely TATA box), (ii) [CT][CT]AACA (positive SHAP value, likely 3’ end of LTR), and (iii) CAACAT[GT]G (negative SHAP value, unknown origin). A complete list of evaluated k-mers in the DNABERT model and their SHAP values is provided as Supplementary File 4. Visualization of signal alignment was conducted in the same way as previously described in section 2.2.1.3 to show the sequence regions containing significant k-mers (Figure 4b).

#### 2.2.2 Superfamily

Models trained to recognize superfamilies were analyzed analogically to those detecting LTRs (subsection 2.2.1.1) and the respective visualizations can be seen in Supplementary Figure 11,12. Interestingly, gProfiler analysis of top 20 TFBS instrumental in superfamily classification by the GBC model did not show any biological process outside general transcriptional regulation as overrepresented in the set. The top 20 list shared 6 TFBS with a similar list from the LTR classification task, namely AHL20, DOF5.3, DOF5.7, ARF 16, ERF069, AT3G46070. Unlike the LTR classification task above, DeepExplainer visualization has not demonstrated the importance of TATA box sequences for superfamily classification, although a few sequences did show some higher SHAP values in the vicinity of the TATA box, some of which could be other core promoter elements preceding TATA. The extreme ends of the LTR sequences remained informative (Supplementary Figure 12), suggesting that these sequences may exist in superfamily variants, or at least are more conserved in one superfamily compared to the other (further analyzed in the following subsection and Table 2).

**Table 2.**
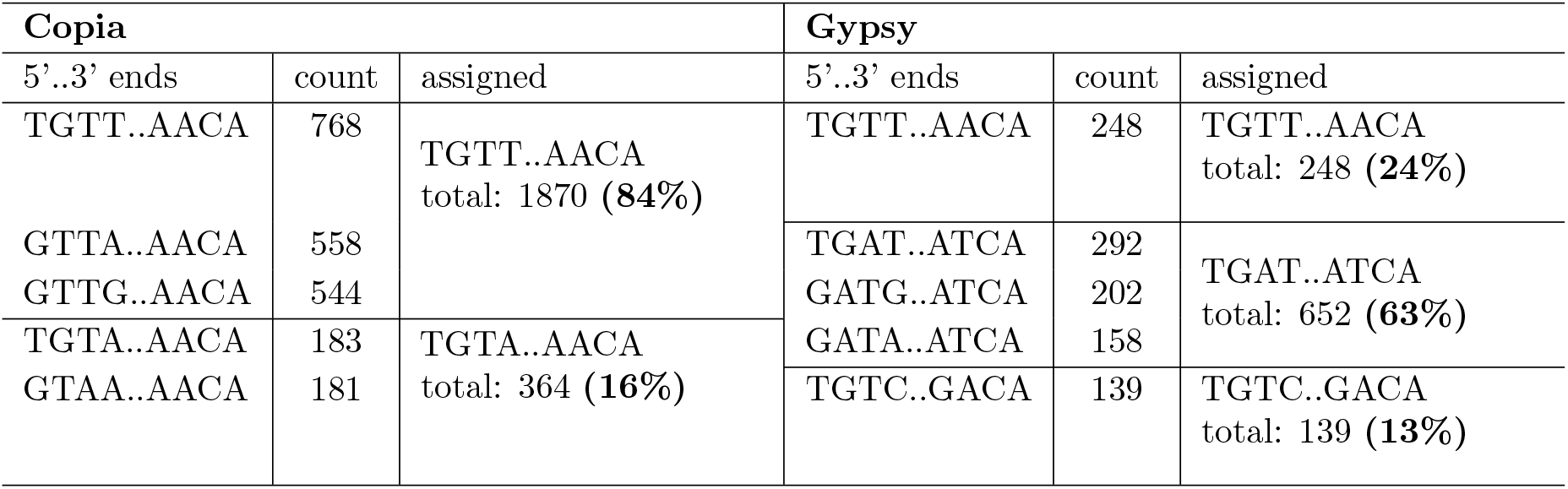
The most represented k-mers at the 5’ and 3’ ends of LTRs. **Left** - Copia superfamily, the most frequent tetramers are 5’-TGTT..AACA-3’; **Right** - Gypsy superfamily, the most frequent tetramers are 5’-TGAT..ATCA-3’.

##### LTR 5’ and 3’ edge analysis

The extremes of LTR sequences (at both the 5’ and 3’ ends) repeatedly surfaced in our explainability analysis results as informative. To get a more detailed picture of sequences present at these locations in various LTR subsets, we also counted the 5’ and 3’ k-mers in the input data. Table 2 shows the assignment of the top five tetramer pairs of Gypsy and Copia superfamily LTR ends to one of the two most frequent pairs, 5’-TGTT..AACA-3’ and 5’-TGAT..ATCA-3’. The assignments were based on the observation that in some LTR annotations, tetramers were apparently shifted by 1 base, presumably because of annotation imprecisions in the sourced files (based on this logic, as an example, GTTA..AACA was assumed to be shifted by 1 base at the 5’-end and therefore assigned to TGTT..AACA). The assumption of a shift in Table 2 was made in all cases where shifting the tetramer by 1bp improved the complementarity of the 5’ and 3’ tetramers and led to the presence of the canonical TG..CA pair. Interestingly, 16 % of Copia edges had only 3bp complementarity, compared to the rest of the five most occuring tetramer pairs in each superfamily shown in the table.

#### 2.2.3 Family

Models trained to recognize families were analyzed analogically to those detecting LTRs (subsection 2.2.1) and the respective visualizations can be viewed in Supplementary Figure 13. Significant signals typical for specific positions within the LTR all but disappeared in visualization of SHAP values from CNN and DNABERT models for individual families (Supplementary File 5). Top TFBS from GBS SHAP analysis had no overlap with the corresponding LTR and superfamily sets. However, when subjected to overrepresentation analysis with gProfiler (Supplementary Figure 10), six of the top 20 TF involved belonged to a functional group responding to plant hormones, specifically auxin and abscisic acid. Related overrepresented biological functions in eight TFBS included cell communication, meristem localization and phyllotaxis (PHY3, AIL6, AIL7, WRKY62, FUS3, ABF2, ABF3, BHLH112).

## 3 Methods

### 3.1 Sequence data

#### 3.1.1 LTR input data

Annotated full-length transposable elements were obtained from Zhou et al. (2021). Available annotations were searched for LTR pairs (a pair of 5’- and 3’-LTRs belonging to the same full-length TE). Element insertion time was estimated based on LTR pair divergence as previously described (Jedlicka et al. 2020). A corresponding FASTA file containing all the sequences further used here is provided as Supplementary File 6.

A separate set of “LTR-negative” sequences was prepared for model training (Supplementary File 7). In supervised learning (used here) a set of DNA sequences that do not contain LTRs is necessary to allow the models to identify classification features that are typical of one set, or the other. Selecting a reasonable negative set was an important and considered step. Apart from generating random sequences, we also strove to include naturally occurring sequences, and sequences with more intricate internal structure. For this, sequences were extracted from the same species as those used for LTR extraction, however this time the annotations were used to avoid regions marked as LTRs. To further the complexity of the training dataset and reduce the influence of easily distinguishable features, a set of sequences generated using Markov chains trained on clusters of LTR sequences was added. First, LTR sequences were clustered using the program CD-HIT (Li and Godzik 2006) with a relatively low similarity threshold of 70 % in order to create larger clusters of less similar sequences. On each cluster, a Markov chain model of order 2 was trained and used to generate artificial sequences (Youens-Clark 2021). These contained 3-mers often found in LTRs, but lacked the spatial and organizational properties of LTR sequences. Counts of these different types of non-LTR sequences are given in Supplementary File 8.

Although training on sequences with more complex differences than those between LTRs and random sequences is a more challenging task, the resulting data provides a more informative trained model, avoiding fitting on trivial or non-biological features, such as sequence composition or features that are typical of any plant genomic sequence.

#### 3.1.2 Cleaning and filtering of input data

Due to annotations and their corresponding reference genomes not always being reliably identified from the published data, only annotations that produced a single-mode distribution of ages were used. We required an LTR sequence alignment identity of 0.7-1.0 which provided 513663 LTR pairs from 79 plant species.

In order to filter the database of LTR sequences and remove redundant, highly identical training examples, a clustering technique based on sequence similarity was applied using the program CD-HIT (Li and Godzik 2006). This approach was tested for sequence similarity *>*95 % and *>*85 % (these numbers were chosen to provide sequence sets of two sizes, while even the smaller set would still contain multiple members of individual TE families, which must have less than 80 % divergence along more than 80 % length by definition (Wicker et al. 2007)). The identity percentage represents the lower boundary of similarity, above which, sequences have been clustered together. One representative was selected from each cluster to be used for training. Applying the 85 % boundary results in a relatively smaller, more strict training database, which should contribute to training more robust models. The resulting LTR database consisted of 176917 sequences and the LTR-negative database of 543310 sequences from 75 species. Supplementary File 9 provides a list of species and the respective LTR counts.

As the original annotated set contained only full-length transposable elements and their corresponding LTR sequences, we wanted to verify that no solo-LTRs from the annotated organisms’ DNA had ended up in the negative training set during the extraction process. Solo-LTR sequences are LTR sequences orphaned through a process of unequal homologous recombination between LTRs of the same full-length element (Vitte and Panaud 2003). The negative training database was therefore aligned to our LTR database in order to filter out any potential solo-LTR candidates.

#### 3.1.3 LTR sequences comparison - classical approaches

Using a custom python script the Jaccard similarity index was counted for k-mers (k = 6) of all the LTRs and/or non-LTRs sequences of the corresponding plant species in order to compare (i) k-mers between Ty1/Copia and Ty3/Gypsy (Supplementary Figure 1a,b); and (ii) LTR and non-LTR sequences originated from the same plan genome (Supplementary Figure 1c). Dendrograms and barplots were generated in R (version 4.3.3) using the default ‘hclust’ function with ‘ape’ package and ggplot2 package, respectively.

### 3.2 Input sequence data preprocessing

Three sequence preprocessing strategies have been implemented, each corresponding to a particular model type and the type of input features that it can handle. These include the identification of transcription factor binding sites, one-hot encodings and k-mer tokenization.

#### 3.2.1 TF binding site identification

To transform the sequences into a fixed-dimension feature space that can be utilized with most conventional models, the sequences were parsed with position weight matrices (PWM) of common plant transcription factor binding sites obtained from the JASPAR database (Castro-Mondragon et al. 2022) (Supplementary File 10). Sequences were searched for TFBS motifs using the Motifs module provided by the biopython package (Cock et al. 2009) and a feature vector of size 656 containing the number of occurrences of each motif was obtained for every input sequence. The vector was then transformed using the term frequency-inverse document frequency (TF-IDF) measure (Manning et al. 2008) to reduce the impact of less specific and common TF binding sites during classification. The advantage of using the TFBS identification approach for training models is the fixed size of the input vector, the easy interpretability and ability to use with most conventional machine learning models. The downside is that we make preemptive assumptions about the data and may omit other important properties, which are hidden within the sequences.

#### 3.2.2 One-hot encoding

In order to maintain the structural information of input sequences, two further approaches to transforming the sequences were undertaken. The first method was transforming the sequences to one-hot encoded vectors. In contrast to other popular methods for encoding DNA sequences, such as k-mer frequency, one-hot encoding allows the CNN filters to fit on specific sequence patterns, maintains sequence structure, and improves the interpretability of the network where each filter can also be viewed as a PWM (Koo and Eddy 2019).

#### 3.2.3 k-mer tokenization

The method used in combination with DNABERT-based fine-tuning tasks was k-mer tokenization. In this case, three different values of k (4, 5, 6) were tested for each classification task. The input sequence was split into k-mers and encoded into numerical vectors using the assigned tokenizer of the pre-trained DNABERT (Ji et al. 2021) for further training in the transformer model.

### 3.3 Models

#### 3.3.1 Gradient Boosting Classifier

We implemented and executed a Gradient Boosting classifier (GBC) pipeline that consisted of three steps. First, we identified TFBS occurrences in input sequences using the src/utils/run jaspar parser.py script. We then used these TFBS occurrences as input for the pipeline containing a TF-IDF transformer and the GBC model, both from the scikit-learn version 1.3.0 (Pedregosa et al. 2011). The corresponding pipelines containing trained models are provided in Supplementary File 11.

#### 3.3.2 CNN-LSTM

We trained the CNN-LSTM model by first preprocessing the sequences using one-hot encoding. We removed unknown bases from the input sequences (represented by “N”) and then encoded each of the 4 bases (A, C, G, T) using a one-hot vector where 1 represents the presence of the particular base in the specified position and all other positions are set to 0. The preprocessing utils are located in the src/utils/CNN utils.py. We trained the model for input sequences of size 4000bp, using the pad sequences function from the keras.utils module version 2.14.0 (Chollet and et al. 2015) to pad sequences shorter than 4000bp with a 0-vector and truncate sequences that are longer. We set up the CNN-LSTM model with optimal parameters detected during the hyperparameter sweep (described in section 3.4.2). The trained models are are provided in Supplementary File 11. Supplementary Figure 14 shows top 20 TFBS identified for each task, overall topology of the model is shown in Supplementary Figure 15.

#### 3.3.3 DNABERT

For the pre-trained DNABERT model fine-tuning process, we loaded the model along with the assigned tokenizer at zhihan1996/DNA bert 6 from the Hugging Face hub (`huggingface.co`) using the transformers module (Wolf et al. 2020)with the AutoTokenizer and AutoModelForMaskedLM functions provided. The fine-tuned models for the specific classification tasks to draw predictions from pooled embeddings are available in Supplementary File 11 and work out-of the box for sequences below 512bps. For sequences above this length, the pooling strategy (described in 3.4.3) needs to be implemented. The functionality for this is provided by the src/utils/seq to embedding.py script. The overall topology of the model is shown in Supplementary Figure 16.

### 3.4 Model training

Training and predictions on CNN-LSTM and DNABERT models were run using a NVIDIA A100 80 GB PCIe GPU. The datasets were divided into training, validation, and testing with the ratio of 70 %, 10 %, and 20 %, respectively.

#### 3.4.1 TFBS models

The TFBS models were subject to a 5-fold cross-validation grid search in order to detect the optimal model and its hyper-parameters (Supplementary Table 1). During this grid search, Random forest classifier, Gradient Boosting classifier and Multilayer Perceptron classifiers were tested for different combinations of parameters (Supplementary File 12). The LTR and superfamily classification models were trained maximizing the binary cross entropy function. The family classifiers were trained maximizing the categorical cross entropy function weighted by the proportion of representatives per class (Supplementary Table 2).

#### 3.4.2 CNN-LSTM

The CNN-LSTM contains an input layer of size 4000, followed by a 1D convolutional layer, a max pooling layer followed by an LSTM layer into the output node. A zero-vector padding technique with masking was used for sequences shorter than 4000bp. All classifiers were trained using the Adam optimizer for 15 epochs, with batches of size 64, using the early stopping criterion with a 3 epoch patience on the validation set to prevent overfitting. The LTR and Superfamily classifiers were trained to optimize the binary cross entropy loss function, whereas the family classifier was trained optimizing the categorical cross entropy function. The models were connected to the Weights and Biases interface (https://wandb.ai) to monitor training progress and a hyperparameter sweep was run to detect the best network hyperparameters (Supplementary File 13).

#### 3.4.3 DNABERT

For the training process of the DNABERT model, it was also connected to the W&B interface, and 3 k-mer sizes were tested for the various classification tasks - 4, 5, 6. All models were trained using the AdamW optimizer for 5 epochs, utilizing the early stopping criterion with a 2 epoch patience on the validation set, to prevent overfitting. The LTR and superfamily classifiers were trained optimizing the binary cross entropy loss function, whereas the family classifier was trained optimizing the BCE with logits loss function (Paszke et al. 2019). These models were trained on sequences under 510 bps in length, 512 being the standard maximum input sequence length of the BERT transformer model. For sequences larger than 510 bps, a window pooling approach was taken, where a window of size 510 corresponding to the input size of the trained model was moved along the input sequence with a stride size of 170 (one third of the of the model’s input length). The classification head of the model was removed, and the produced embedding vector of size 768 was average-pooled along the sequence, generating a final vector of size 768 containing averaged embeddings along the sequence. A convolutional network model was then trained to classify sequences based on the pooled embedding vector. This network uses 32 filters of size 3 pooled into a dense layer of size 32 and a logistic sigmoid at the activation function in the output layer for the LTR and superfamily classification tasks and a softmax layer of size 15 for the family classification task.

### 3.5 Trained model interpretation

#### 3.5.1 SHAP

The SHAP (SHapley Additive exPlanations) algorithm has its roots in cooperative game theory. It is a model-agnostic approach used for estimating the impact that input features have on the output of the model. Shapley values exhibit desirable properties such as efficiency, symmetry, and additivity, making them an ideal foundation for understanding the contribution of each feature to a given prediction. Due to the additive nature of Shapley values, they may be used for local, instance-wise explanation, as well as global understanding of input features across multiple instances when aggregated.

The algorithm works by training a model *f*_*S*∪*i*_ with feature i present during the training and a model *f*_*S*_ with feature i not present. The outputs of these models are then compared and the SHAP value for feature i is computed as the difference of outputs of the models *f*_*S*∪*i*_ and *f*_*S*_ scaled by the weighted average of all possible differences.

To produce the SHAP values used in this study, the python module SHAP (version 0.44.1) was used. The gradient boosting classifier feature importance was interpreted using the TreeExplainer (Lundberg et al. 2020) class provided by the package on the full testing dataset of LTRs and LTR-negatives in the case of LTR classification, and the full set of only LTRs in the case of superfamily classification.

For interpreting the CNN-LSTM hybrid network model, the LTR test set was subsampled down to 2000 instances, and was parsed using the DeepExplainer module of the SHAP package to explain feature importance of input sequence positions.

For interpreting the k-mer based fine-tuned DNABERT model, the Explainer module with automatic selection of estimator was used to interpret the importance of k-mers in sequences within 512 length, for the selected set of 2000 LTR sequences. In order to obtain the importance of particular k-mers, we aggregated the SHAP values for each k-mer across the selected subsample. Their corresponding values were then scaled using Min-Max scaling separately for k-mers with largest negative and largest positive contributions.

Additionally, we analyzed the importance of specific regions of LTRs (sequence start, TATA box, TSS, and sequence end). First, we predicted the positions of TATA and TSS sites using TSSPlant (Shahmuradov et al. 2017). Next, we assessed the importance of each sequence position across 2000 subsampled LTR sequences by either calculating the sum of squares of SHAP values in each position (CNN-LSTM model) or calculating the mean SHAP value of all k-mers containing a given position (BERT). Median position importance, centered around specific regions, was then visualized using the plotHeatmap function from deepTools (Ramírez et al. 2016).

#### 3.5.2 CNN filter analysis

To analyze the learned CNN filters, they were first extracted from the trained network, then normalized in the following way: 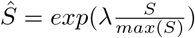 where Ŝ represents the normalized filter, S the original filter and *λ* is a scaling factor whose value was set to 3 as suggested in Koo and Eddy (2019). These normalized filters were then converted to the MEME format using the jaspar2meme tool from the meme suite version 5.5.5. (Bailey et al. 2015) and compared to the JASPAR CORE 2022 (Castro-Mondragon et al. 2022) plant database using the Tomtom motif comparison tool (Bailey et al. 2015) with a cutoff E-value of 0.1.

## 4 Discussion

We used machine learning methods to predict long terminal repeats (LTR) of plant LTR retrotransposons and to classify LTR sequences into retrotransposon families. Our results show that the models used here recognized biologically relevant motifs, such as core promoter elements (TATA box), as well as development- and stress-related subclasses of TF binding sites. Our analysis also reinforced the importance of 5’- and 3’-edges in LTR identity.

While our work is not the first to apply machine learning methods to LTR retrotransposon analysis, none of the previous studies analyzed LTRs in isolation as we did here. One of the earliest ML approaches, based on full-length TE sequences, was reported by Schietgat et al. (2018) who used Random Forest-based models to detect and classify LTR retrotransposons into superfamilies, achieving average F1 values of 0.56. Orozco-Arias et al. (2021) trained a multi-layer perceptron model based on k-mers to classify full-length LTR TEs. The F1 scores on their data reached 0.95. This is a better performance than our deep learning models here at or above the superfamily level (0.73-0.85; Table 1), however it is also expected, since the internal parts of LTR TEs contain protein coding sequences that are more amenable to sequence alignment, as such form the basis of TE classification systems, and are generally easier to detect and cluster. Other previous works aimed at classifying LTR TEs as a class among other repeat classes used neural networks and hierarchical repeat sequence clustering (Abrusán et al. 2009, Nakano et al. 2017) to achieve precision of LTR-TE classification 0.88 and 0.94. These variances can be ascribed to different motivation. While we focused on model explainability and the associated detection of biological motifs in the analyzed LTR sequences, the other studies were mostly motivated by increasing the speed and/or precision of the classification tasks compared to possibly simpler but time-consuming procedures, such as sequence alignment. The use of isolated LTRs allowed us to focus on specific sequences that typically make up regulatory DNA, not only in LTRs but also in promoters and enhancers. Unlike the above approaches, we also carried out classification at the family level. While these models were the most difficult to analyze for explainability, and the least informative compared to the two higher levels, they still achieved a respectable F1 of 0.68-0.75.

Looking at previous attempts in this area, clearly models with a CNN component tend to be the most popular and give the best results (da Cruz et al. 2021, Yan et al. 2020); see also Table 1).

We tested three different techniques to achieve model explainability and identify features that the models “learned”, which then contributed most to model accuracy. Interpretation of DNABERT attention heads (not shown) was problematic. Among other things, we did not find an effective way to correlate the data with the other methods (CNN filter analysis and SHAP) and therefore decided not to pursue this avenue of investigation. CNN filter analysis has shown that many of the filters learned in the neural network had resemblance to known JASPAR TFBS motifs and served to pinpoint the most prominent TFBS recognized by the models. Their biological underpinnings are discussed below. It turned out that SHAP was the most effective method to analyze the trained models, which allowed us to identify specific sequence motifs used by the models, such as the TATA-box motif and 5’- and 3’- ends of LTRs, that contributed most to LTR identification and classification and are identical to motifs described in plant TEs before (Rocheta et al. 2012). Also, being a model-agnostic method, the use of SHAP allowed us to compare influential features across models using the same metric.

The dependence of models on TFBSs in LTRs is consistent with the concept that LTRs are regulatory regions capable of controlling the transcription of elements in a spatially and temporally specific manner (Wicker et al. 2007). By searching biological roles of the most prominent TFBSs, we found them to be associated particularly with (i) transcriptional activation of genes in stress conditions (DREB1, REF6, ERF7, ARR1), (ii) binding sites for transcription factors (TFs) acting during flowering and germline development (RAMOSA1, CDF5, DOF5.3, E2FA,C,D,E, MYB24, NID1,TB1, AT3G46070), (iii) binding sites for tissue-specific transcriptional repressors (AHL20, CDF2, ARF2), and (iv) binding sites for chromatin remodelers involved in DNA demethylation (REF6). This observation gained using LTRs from 75 plant species here should be interpreted with caution because TFs form large gene families of neo-functionalized and sub-functionalized genes sharing identical or similar TFBSs. TFs typically account for 5-10 % of genes in a species genome (Yuan et al. 2024), for example, Arabidopsis thaliana has approximately 2300 TFs, which corresponds to 8.3 % of its total genes (Hong 2016). Some TFs also either require the binding of homodimers to two TFBSs at some distance apart or interaction with other TFs bound to a given locus to initiate transcription or other processes (Boer et al. 2014, Strader et al. 2022). Moreover, the roles of individual TFs have only been studied in a few model species to date, and it is unclear to what extent their functions are conserved in plants.

However, some of the prominent TFBSs recognized by the models have already been found and functionally validated in TEs. Therefore, we assume that our models have used TFBSs preferred in LTRs and related to general rules for transcriptional regulation of LTR retrotransposons. Of particular importance are TFBSs for binding stress-response TFs. Activation of TEs by abiotic and biotic stress is supported by a wealth of experimental data in A. thaliana (e.g. (Duan et al. 2008, Matsunaga et al. 2011), rice (Jiao and Deng 2007), sunflower (Mascagni et al. 2020) and other species (Ito 2022). Although earlier studies linked the activation of TEs by stress to epigenetic changes (euchromatinization of TEs), a number of TEs are now known to contain TFBSs identical to those of stress-responsive genes. A textbook example is ONSEN, a heat-induced (high temperature induced) LTR retrotransposon containing a heat shock element (HSE) for heat shock factor binding (Cavrak et al. 2014, Ito et al. 2011). In maize most TEs (gypsy, copia, LINEs) activated by stress contain motifs for stress-responsive DREB/CBF transcription factors (Makarevitch et al. 2015), which have been recognized by our ML models. DREB/CBF and REF6 TFBSs have been detected in Gypsy and Copia TEs activated by heat stress in Arabidopsis (Deneweth et al. 2022). REF6 is a plant-unique H3K27 demethylase that targets DNA motifs via its zinc-finger (ZnF) domain (Lu et al. 2011). Its presence in TEs suggests that TEs are able to actively resist the host methylation machinery and/or control their epigenetic state in response to stress conditions. In addition, there is increasing evidence that TEs, by transferring stress-response TFBSs to the vicinity of genes, rewire new transcriptional networks that enable the host adaptation to stress (Deneweth et al. 2022, Hénaff et al. 2014, Qiu and Köhler 2020) and changing environmental conditions (Quadrana 2020).

Another frequent group of TFBSs is bound by TFs expressed in floral meristems and reproductive organs. TB1 (identified here by ML models) has been previously confirmed in the Hopscotch retrotransposon in maize, where it is expressed in developing glumes (Dong et al. 2019). E2F TFBSs were found in several families of TEs in Brassica species, and E2Fa binding to TEs has been functionally validated in vivo (Hénaff et al. 2014). E2F TFs regulate various processes mostly in developing pistils and anthers, and frequently TE-harbored MYB24, NID1, CDF5, and AT3G46070 TFs also show localized expression in stamens (based on https://bar.utoronto.ca/efp/cgi-bin/efpWeb.cgi, Klepikova Atlas). These findings suggest that TEs prefer certain short-term and localized windows in their host’s life cycle for transcription and transposition. Transposition in floral meristems or in reproductive cells allows TEs to minimize their spatiotemporal activity, thereby lowering the risk of reducing host fitness by deleterious insertions in somatic cells, while increasing the probability of transmitting new TE copies to the next generation. Clues to this behavior can be seen in dioecious plants with heteromorphic sex chromosomes. For example, in Silene latifolia and Rumex acetosa, the accumulation/absence of most LTR retrotransposons on Y chromosomes can be explained by transposition in either the male or female reproductive organs (Cermak et al. 2008, Filatov et al. 2009, Hobza et al. 2017, Jesionek et al. 2021, Kubat et al. 2014, Steflova et al. 2013). A very similar situation emerges in animals as well. For example, many TEs harbor DNA binding sites for pluripotency factors and are transiently expressed during the embryonic genome activation of primates (Pontis et al. 2022).

Taken together, the ML tools opted for TFBSs, many of which have been independently described by other methods. We consider them to be indicative of the biology of TEs and the TE/host interaction. We can speculate that, in general, LTRs contain more often binding sites of TFs that ensure reproductive cell-specific activity or activity triggered by biotic and abiotic stresses. This is advantageous for both TEs and the host because (i) host viability is not threatened by deleterious TEs in somatic cells, (ii) transgenerational reproduction of TEs is ensured, and (iii) the evolutionary plasticity of the host genome is increased by new regulatory networks (Gebrie 2023). On the other hand, no single TFBS defines a specific taxonomic group of TEs suggesting that TEs can co opt new TFs from other TEs and genes, and adapt their strategy to changing conditions in the host genome.

Our machine learning approach could be advantageous not only for a better LTR retrotransposon and solo LTR identification and annotation but could be useful also for the prediction of potential TF binding sites within LTR. This way, our tool can also contribute to revealing the involvement of these mobile genetic elements in cellular regulatory networks.

## 5 Conclusion

In this work we tested the ability of deep learning techniques to learn features specific for certain sets of plant LTR sequences, and when combined with explainability analysis, to pinpoint regions of LTRs responsible for their accuracy. We found three features used by the trained models: i) 5’- and 3’- edges, ii) TATA-box region, and iii) TFBS motifs and discussed their biological relevance. Our work shows the applicability of the used models and the associated explainability analysis to the study of regulatory sequences and their classification.

## Supplementary Information

### Supplementary Figures

~~~
Supplementary Figure 1 - LTR sequences comparison - classical approaches.
Supplementary Figure 2 - LTR detection accuracy
Supplementary Figure 3 - LTR detection accuracy
Supplementary Figure 4 - Superfamily classification accuracy
Supplementary Figure 5 - Superfamily classification accuracy
Supplementary Figure 6 - Family classification accuracy
Supplementary Figure 7 - Family classification accuracy
Supplementary Figure 8 - gProfiler GOSt analysis of the top 20 GBS model TFBS
Supplementary Figure 9 - gProfiler GOSt analysis of the top 20 CNN model TFBS
Supplementary Figure 10 - gProfiler GOSt analysis of the top 20 CNN model TFBS
Supplementary Figure 11 - Main results of explainability analysis - superfamily
Supplementary Figure 12 - DeepExplainer analysis of trained superfamily detection models
Supplementary Figure 13 - Main results of explainability analysis - family
Supplementary Figure 14 - Top TF binding sites in CNN filter analysis
Supplementary Figure 15 - CNN-LSTM model topology
Supplementary Figure 16 - DNABERT model topology.
~~~

### Supplementary Figures

~~~
Supplementary Table 1 - Tested hyperparameters
Supplementary Table 2 - Loss functions
~~~

### Supplementary Files

~~~
File1_GBC_SHAP_values.tab
File2_CNN_filters.tab
File3_kmer_motifs.pdf
File4_DNABERT_kmer_SHAP_values.tab
File5_CNN_BERT_Grid.pdf
File6_LTR_sequences.fa.gz
File7_LTR-negative_sequences.fa.gz
File8_non-LTR_counts.tab
File9_LTR_species_counts.tab
File10_JASPAR_matrices.tab
File11_Trained_models.zip
File12_Gridsearch_results.json
File13_Hyperparameter_sweep.zip
~~~

## Declarations

### Ethics approval and consent to participate

Not applicable

### Consent for publication

Not applicable

### Availability of data and materials

Supplementary figures, tables, files, frozen source code and data available via Zenodo archive doi://10.5281/zenodo.11555642.

### Authors’ contributions

JH, PJ and ML participated in data collection and preparation, JH carried out model implementation and training; JH, PJ, MK and ML generated final results and visualizations; all authors participated in data interpretation and writing the manuscript.

## Acknowledgments

We thank Christopher Johnson for critical reading of this manuscript.

## Code availability

GIT repository https://github.com/jakubhorvath/LTR_classification

