## Supplemental Figures and Tables for "Detection and classification of long terminal repeat sequences in plant LTR-retrotransposons and their analysis using explainable machine learning"

### Supplementary figures

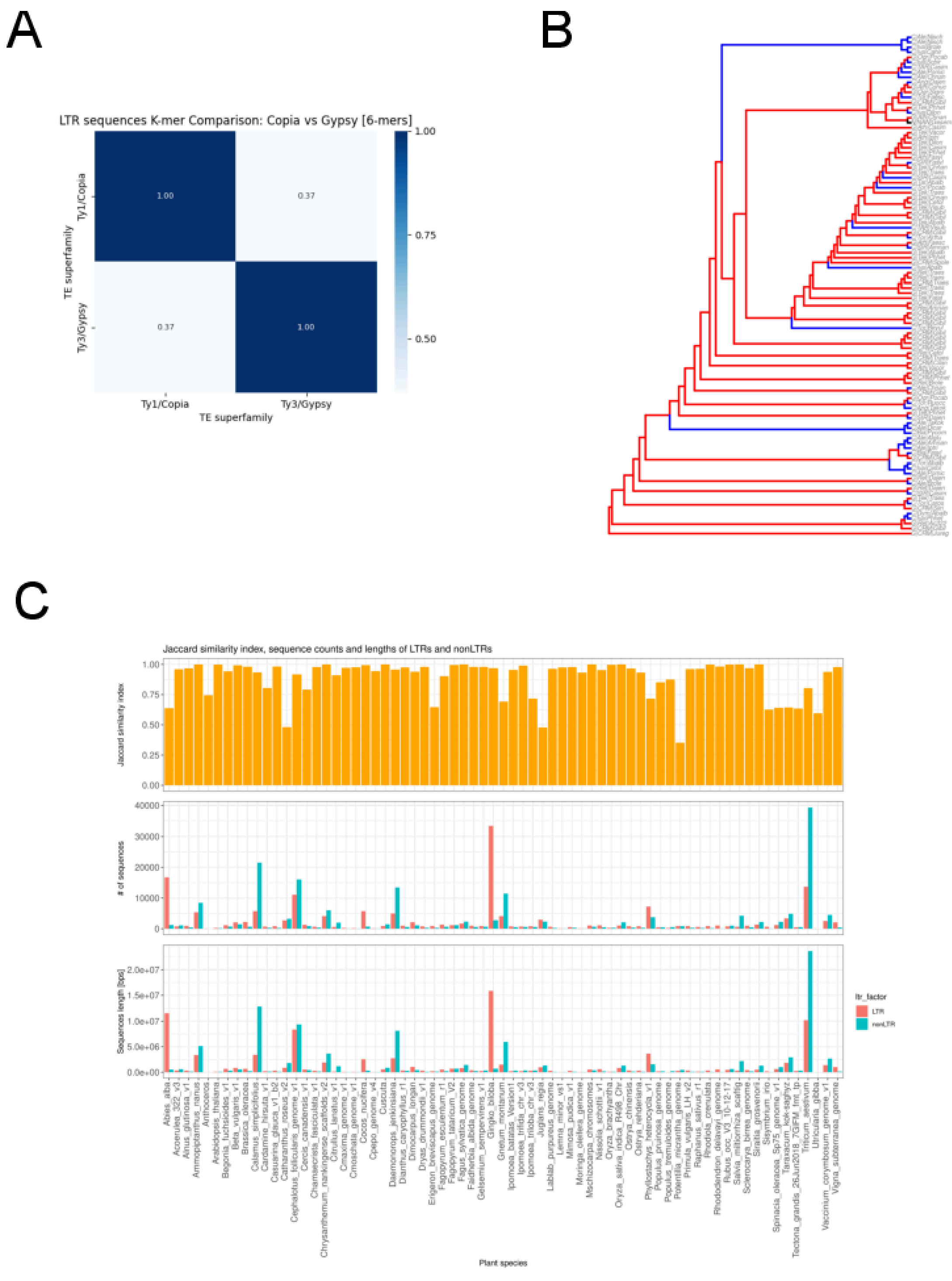


***Supplementary Figure 1 - LTR sequences comparison - classical approaches.*** *Unique k-mers of Ty1/Copia and Ty3/Gypsy LTRs [k = 6; Jaccard Similarity Index (JSI)] (A); Dendrogram constructed from JSI (B); and k-mers occurrence comparison in LTRs and corresponding non-LTR sequences from the genomes of relevant plants as controls (C) .*

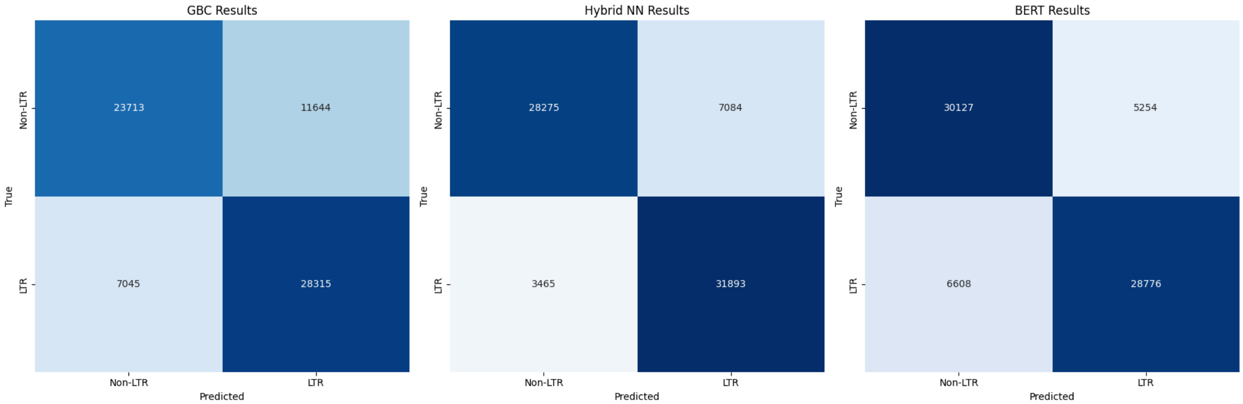


***Supplementary Figure 2 - LTR detection accuracy**. True and false positives/negatives for the three models (GBC, CNN-LSTM and DNABERT) in the LTR binary classification task. The numbers in the contingency table cells are counts of classified LTRs for each combination of attributes.*

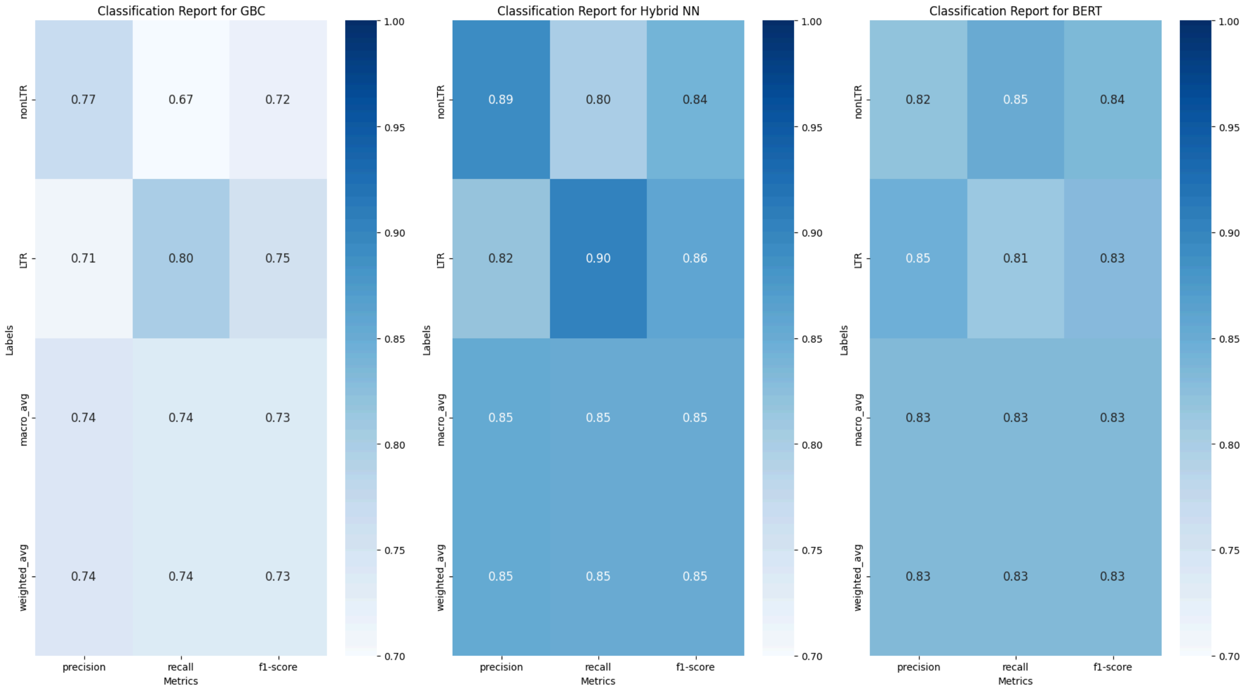


***Supplementary Figure 3 - LTR detection accuracy**. Accuracy characteristics (precision, recall and F1 value) calculated from values in Supplementary Figure 1.*

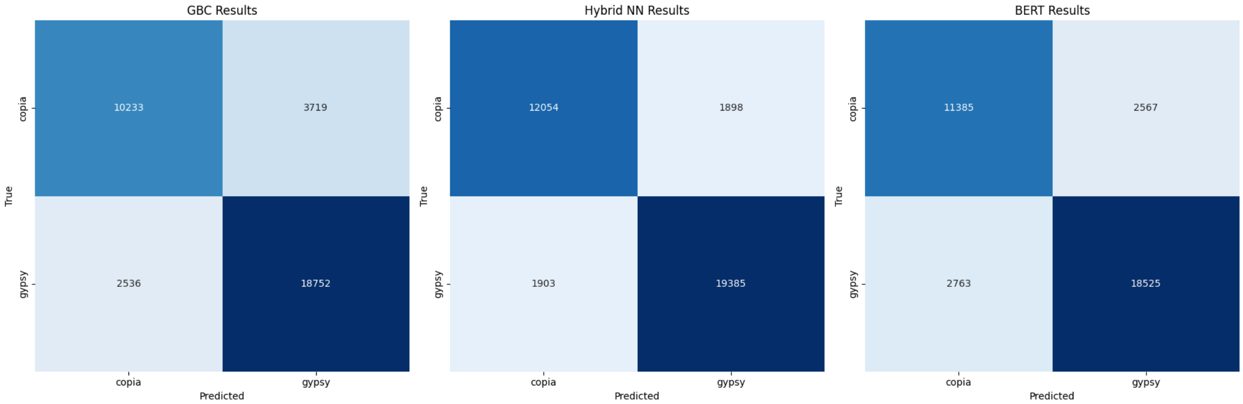


***Supplementary Figure 4 - Superfamily classification accuracy***. *True and false positives/negatives for the three models (GBC, CNN-LSTM and DNABERT) in the LTR binary classification task. The numbers in the contingency table cells are counts of classified LTRs for each combination of attributes.*

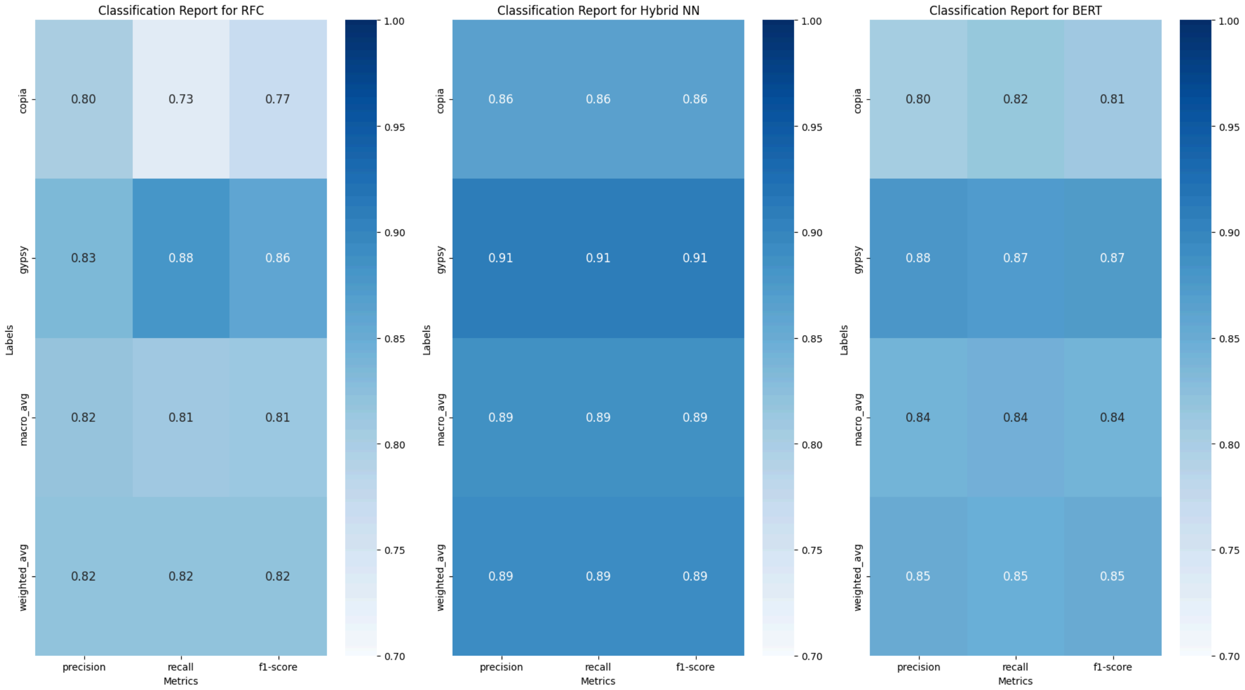


***Supplementary Figure 5 - Superfamily classification accuracy***. *Accuracy characteristics (precision, recall and F1 value) calculated from values in Supplementary Figure 1.*

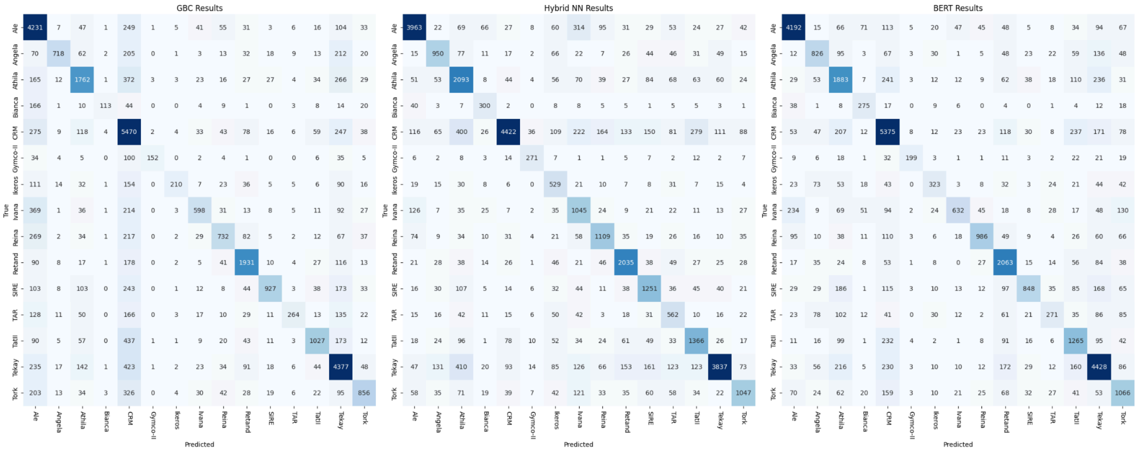


***Supplementary Figure 6 - Family classification accuracy***. *True and false positives/negatives for the three models (GBC, CNN-LSTM and DNABERT) in the LTR binary classification task. The numbers in the contingency table cells are counts of classified LTRs for each combination of attributes.*

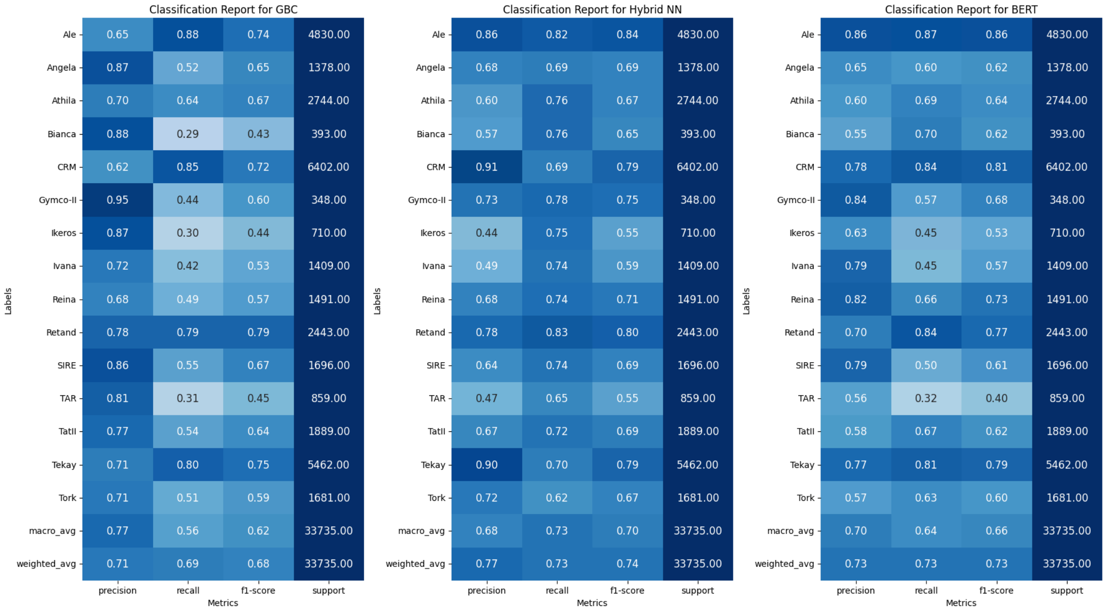


***Supplementary Figure 7 - Family classification accuracy***. *Accuracy characteristics (precision, recall and F1 value) calculated from values in Supplementary Figure 1.*

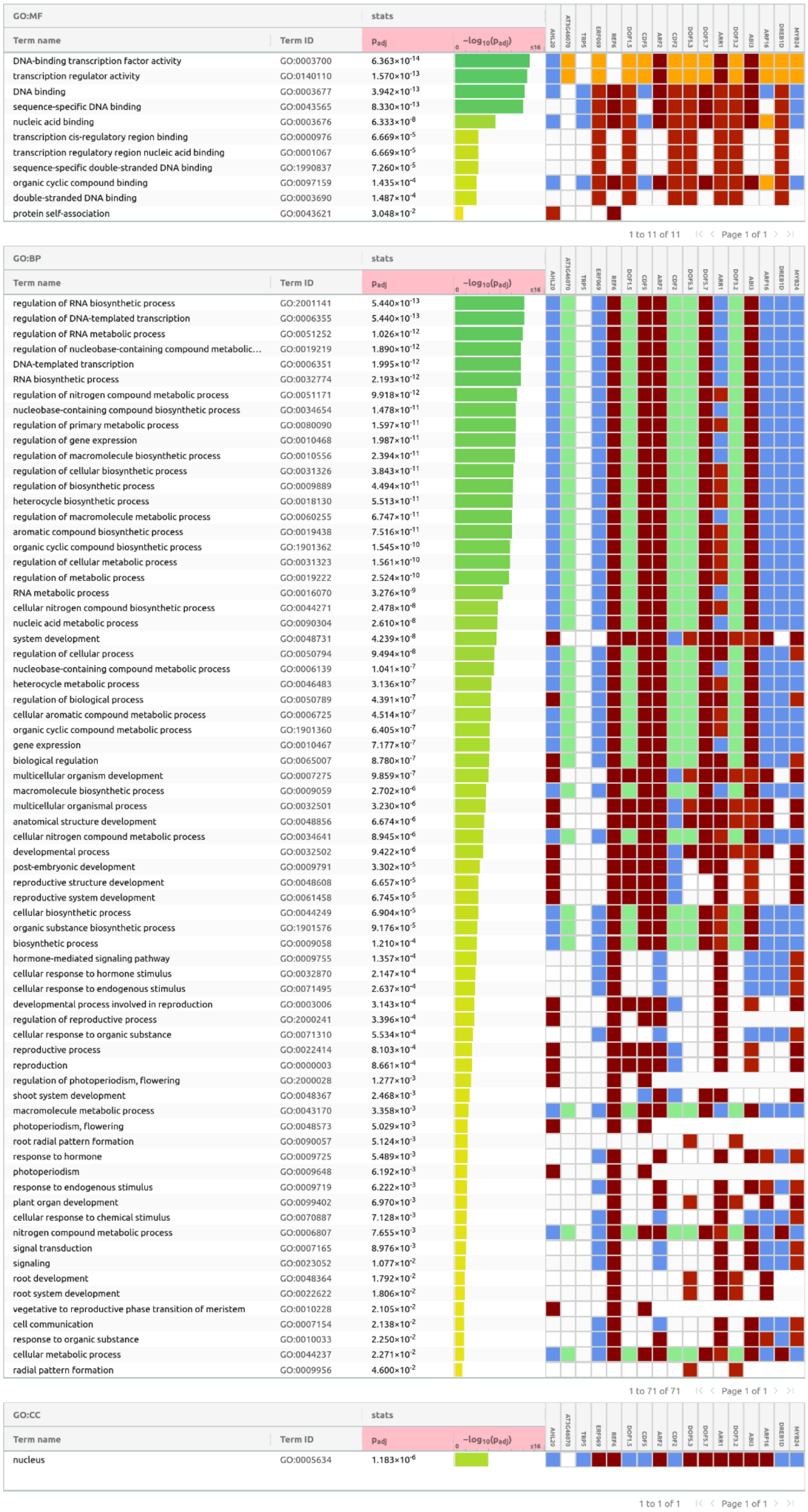


**Supplementary Figure 8 - gProfiler GOSt analysis of the top 20 GBS model TFBS.**

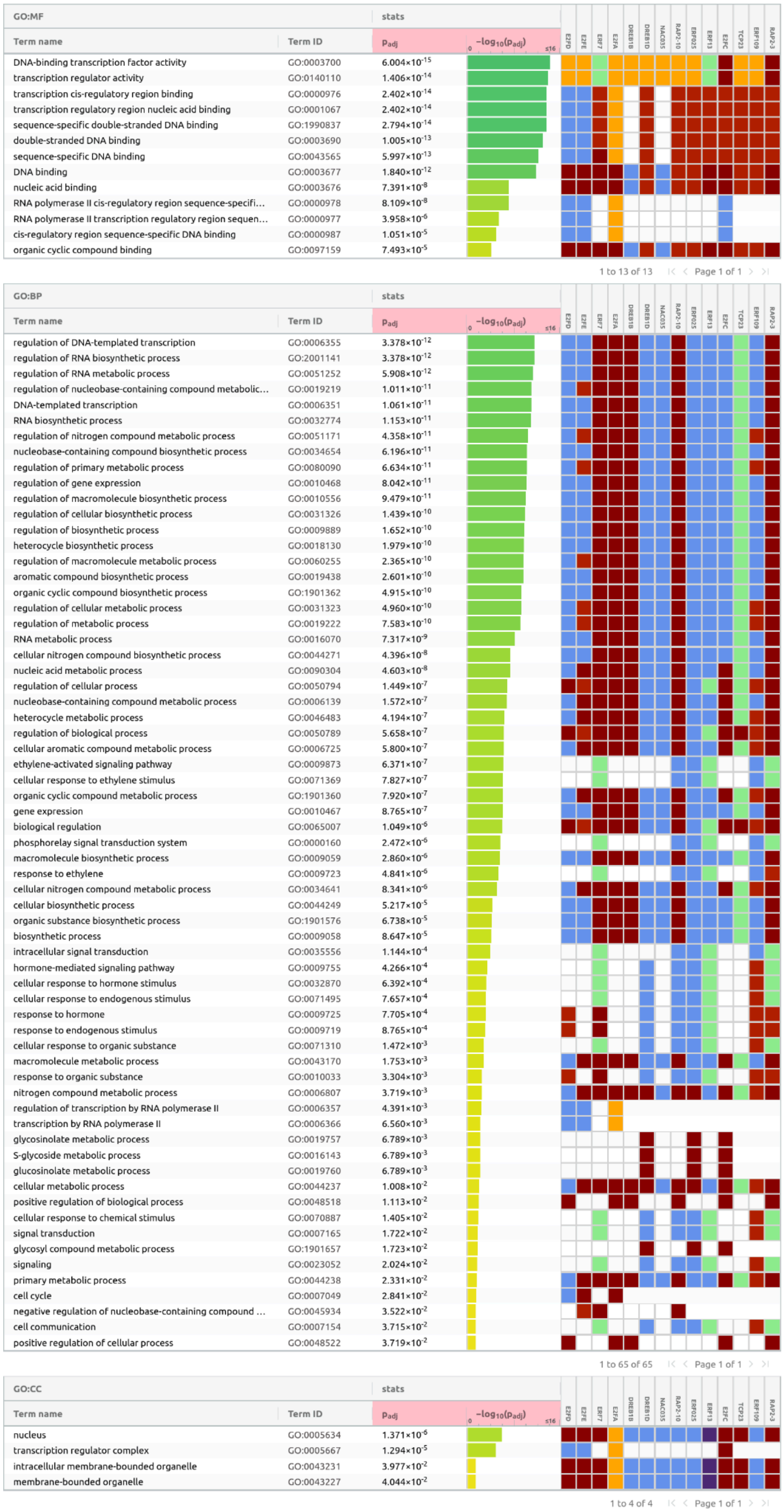


| Term name | Term ID | $p_{adj}$ |
|---|---|---|
| DNA-binding transcription factor activity | GO:0003700 | $6.004\times10^{-15}$ |
| transcription regulator activity | GO:0140110 | $1.406\times10^{-14}$ |
| transcription cis-regulatory region binding | GO:0000976 | $2.402\times10^{-14}$ |
| transcription regulatory region nucleic acid binding | GO:0001067 | $2.402\times10^{-14}$ |
| sequence-specific double-stranded DNA binding | GO:1990837 | $2.794\times10^{-14}$ |
| double-stranded DNA binding | GO:0003690 | $1.005\times10^{-13}$ |
| sequence-specific DNA binding | GO:0043565 | $5.997\times10^{-13}$ |
| DNA binding | GO:0003677 | $1.840\times10^{-12}$ |
| nucleic acid binding | GO:0003676 | $7.391\times10^{-8}$ |
| RNA polymerase II cis-regulatory region sequence-specifi… | GO:0000978 | $8.109\times10^{-8}$ |
| RNA polymerase II transcription regulatory region sequen… | GO:0000977 | $3.958\times10^{-6}$ |
| cis-regulatory region sequence-specific DNA binding | GO:0000987 | $1.051\times10^{-5}$ |
| organic cyclic compound binding | GO:0097159 | $7.493\times10^{-5}$ |

| Term name | Term ID | $p_{adj}$ |
|---|---|---|
| regulation of DNA-templated transcription | GO:0006355 | $3.378\times10^{-12}$ |
| regulation of RNA biosynthetic process | GO:2001141 | $3.378\times10^{-12}$ |
| regulation of RNA metabolic process | GO:0051252 | $5.908\times10^{-12}$ |
| regulation of nucleobase-containing compound metabolic… | GO:0019219 | $1.011\times10^{-11}$ |
| DNA-templated transcription | GO:0006351 | $1.061\times10^{-11}$ |
| RNA biosynthetic process | GO:0032774 | $1.153\times10^{-11}$ |
| regulation of nitrogen compound metabolic process | GO:0051171 | $4.358\times10^{-11}$ |
| nucleobase-containing compound biosynthetic process | GO:0034654 | $6.196\times10^{-11}$ |
| regulation of primary metabolic process | GO:0080090 | $6.634\times10^{-11}$ |
| regulation of gene expression | GO:0010468 | $8.042\times10^{-11}$ |
| regulation of macromolecule biosynthetic process | GO:0010556 | $9.479\times10^{-11}$ |
| regulation of cellular biosynthetic process | GO:0031326 | $1.439\times10^{-10}$ |
| regulation of biosynthetic process | GO:0009889 | $1.652\times10^{-10}$ |
| heterocycle biosynthetic process | GO:0018130 | $1.979\times10^{-10}$ |
| regulation of macromolecule metabolic process | GO:0060255 | $2.365\times10^{-10}$ |
| aromatic compound biosynthetic process | GO:0019438 | $2.601\times10^{-10}$ |
| organic cyclic compound biosynthetic process | GO:1901362 | $4.915\times10^{-10}$ |
| regulation of cellular metabolic process | GO:0031323 | $4.960\times10^{-10}$ |
| regulation of metabolic process | GO:0019222 | $7.583\times10^{-10}$ |
| RNA metabolic process | GO:0016070 | $7.317\times10^{-9}$ |
| cellular nitrogen compound biosynthetic process | GO:0044271 | $4.396\times10^{-8}$ |
| nucleic acid metabolic process | GO:0090304 | $4.603\times10^{-8}$ |
| regulation of cellular process | GO:0050794 | $1.449\times10^{-7}$ |
| nucleobase-containing compound metabolic process | GO:0006139 | $1.572\times10^{-7}$ |
| heterocycle metabolic process | GO:0046483 | $4.194\times10^{-7}$ |
| regulation of biological process | GO:0050789 | $5.658\times10^{-7}$ |
| cellular aromatic compound metabolic process | GO:0006725 | $5.800\times10^{-7}$ |
| ethylene-activated signaling pathway | GO:0009873 | $6.371\times10^{-7}$ |
| cellular response to ethylene stimulus | GO:0071369 | $7.827\times10^{-7}$ |
| organic cyclic compound metabolic process | GO:1901360 | $7.920\times10^{-7}$ |
| gene expression | GO:0010467 | $8.765\times10^{-7}$ |
| biological regulation | GO:0065007 | $1.049\times10^{-6}$ |
| phosphorelay signal transduction system | GO:0000160 | $2.472\times10^{-6}$ |
| macromolecule biosynthetic process | GO:0009059 | $2.860\times10^{-6}$ |
| response to ethylene | GO:0009723 | $4.841\times10^{-6}$ |
| cellular nitrogen compound metabolic process | GO:0034641 | $8.341\times10^{-6}$ |
| cellular biosynthetic process | GO:0044249 | $5.217\times10^{-5}$ |
| organic substance biosynthetic process | GO:1901576 | $6.738\times10^{-5}$ |
| biosynthetic process | GO:0009058 | $8.647\times10^{-5}$ |
| intracellular signal transduction | GO:0035556 | $1.144\times10^{-4}$ |
| hormone-mediated signaling pathway | GO:0009755 | $4.266\times10^{-4}$ |
| cellular response to hormone stimulus | GO:0032870 | $6.392\times10^{-4}$ |
| cellular response to endogenous stimulus | GO:0071495 | $7.657\times10^{-4}$ |
| response to hormone | GO:0009725 | $7.705\times10^{-4}$ |
| response to endogenous stimulus | GO:0009719 | $8.765\times10^{-4}$ |
| cellular response to organic substance | GO:0071310 | $1.472\times10^{-3}$ |
| macromolecule metabolic process | GO:0043170 | $1.753\times10^{-3}$ |
| response to organic substance | GO:0010033 | $3.304\times10^{-3}$ |
| nitrogen compound metabolic process | GO:0006807 | $3.719\times10^{-3}$ |
| regulation of transcription by RNA polymerase II | GO:0006357 | $4.391\times10^{-3}$ |
| transcription by RNA polymerase II | GO:0006366 | $6.560\times10^{-3}$ |
| glycosinolate metabolic process | GO:0019757 | $6.789\times10^{-3}$ |
| S-glycoside metabolic process | GO:0016143 | $6.789\times10^{-3}$ |
| glucosinolate metabolic process | GO:0019760 | $6.789\times10^{-3}$ |
| cellular metabolic process | GO:0044237 | $1.008\times10^{-2}$ |
| positive regulation of biological process | GO:0048518 | $1.113\times10^{-2}$ |
| cellular response to chemical stimulus | GO:0070887 | $1.405\times10^{-2}$ |
| signal transduction | GO:0007165 | $1.722\times10^{-2}$ |
| glycosyl compound metabolic process | GO:1901657 | $1.723\times10^{-2}$ |
| signaling | GO:0023052 | $2.024\times10^{-2}$ |
| primary metabolic process | GO:0044238 | $2.331\times10^{-2}$ |
| cell cycle | GO:0007049 | $2.841\times10^{-2}$ |
| negative regulation of nucleobase-containing compound … | GO:0045934 | $3.522\times10^{-2}$ |
| cell communication | GO:0007154 | $3.715\times10^{-2}$ |
| positive regulation of cellular process | GO:0048522 | $3.719\times10^{-2}$ |

| Term name | Term ID | $p_{adj}$ |
|---|---|---|
| nucleus | GO:0005634 | $1.371\times10^{-6}$ |
| transcription regulator complex | GO:0005667 | $1.294\times10^{-5}$ |
| intracellular membrane-bounded organelle | GO:0043231 | $3.977\times10^{-2}$ |
| membrane-bounded organelle | GO:0043227 | $4.044\times10^{-2}$ |

**Supplementary Figure 9 - gProfiler GOSt analysis of the top 20 CNN model TFBS**

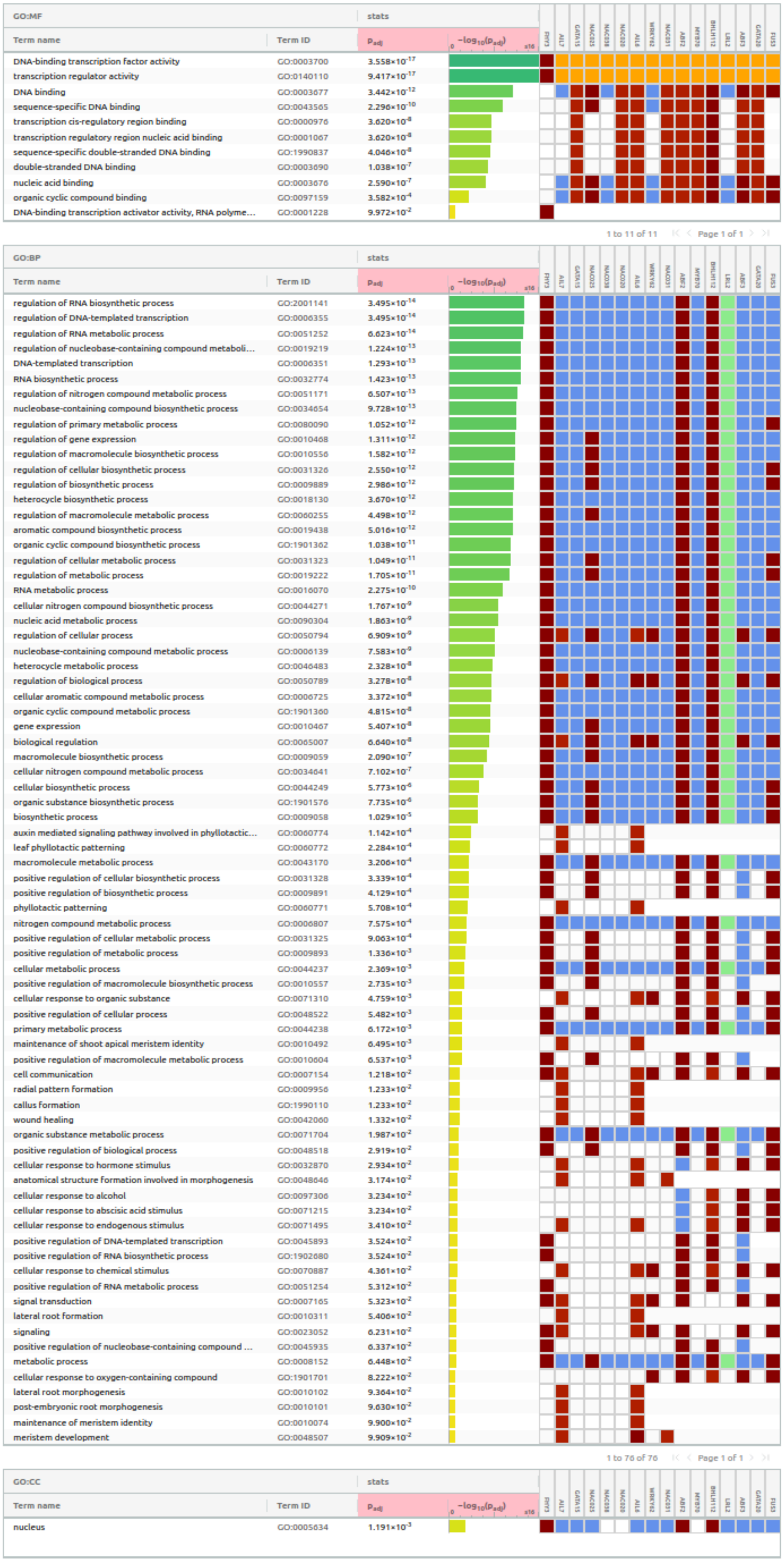


**Supplementary Figure 10 - gProfiler GOSt analysis of the top 20 CNN model TFBS**

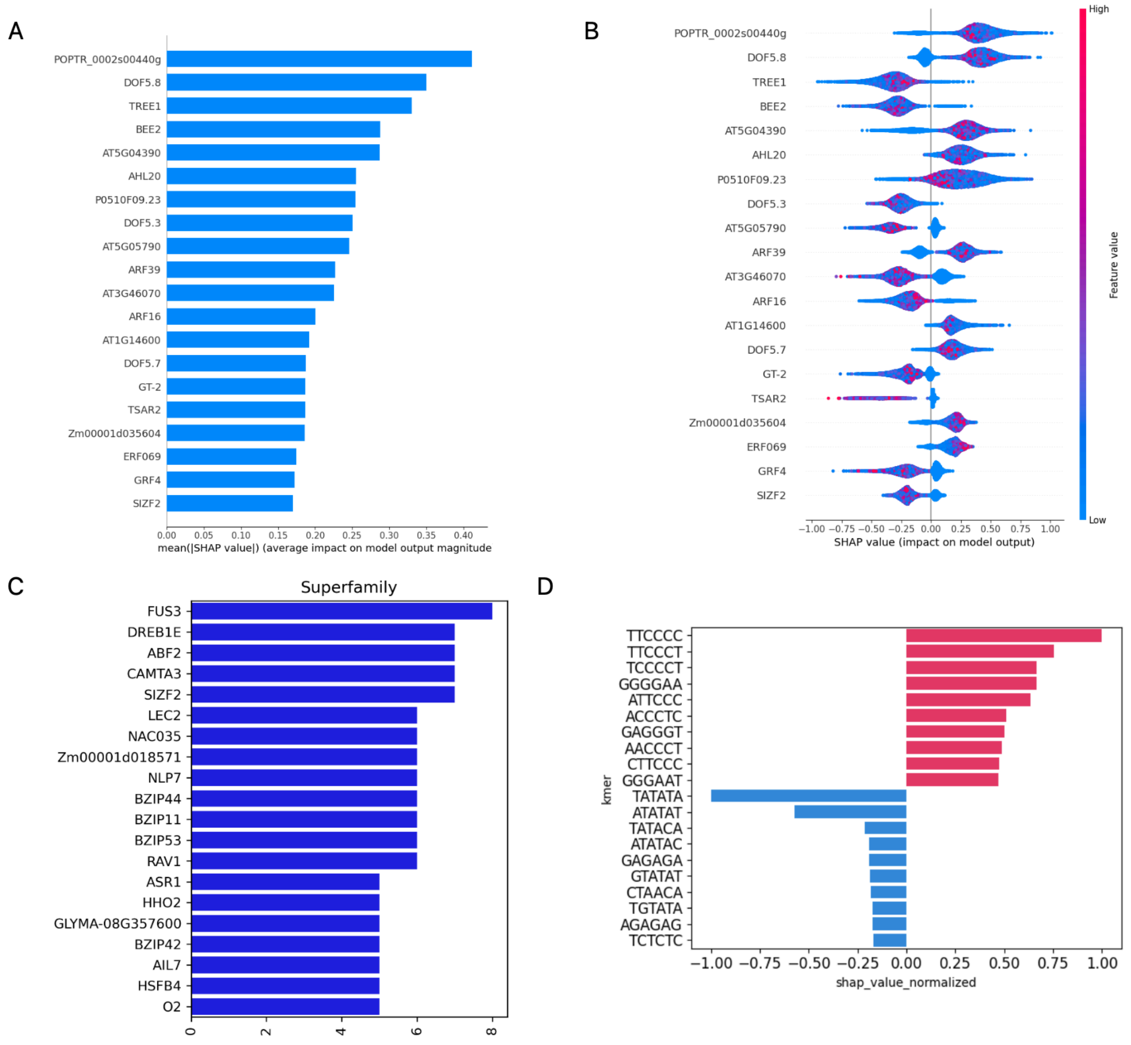


***Supplementary Figure 11 - Main results of explainability analysis carried out on trained models for superfamily classification.*** *A - Mean SHAP contribution of TFBS as input features to the GBC model for LTR classification. B - Beeswarm plot showing extent to which higher/lower values of input features influence model output. Negatively valued contributions are representative of the Copia superfamily, while positive contributions are representative of the Gypsy superfamily C - TomTom hits of first-layer CNN filters on JASPAR Core 2022 database D - Contribution of most significant k-mers used as input features for the DNABERT model. Values identical to those described for* ***B***

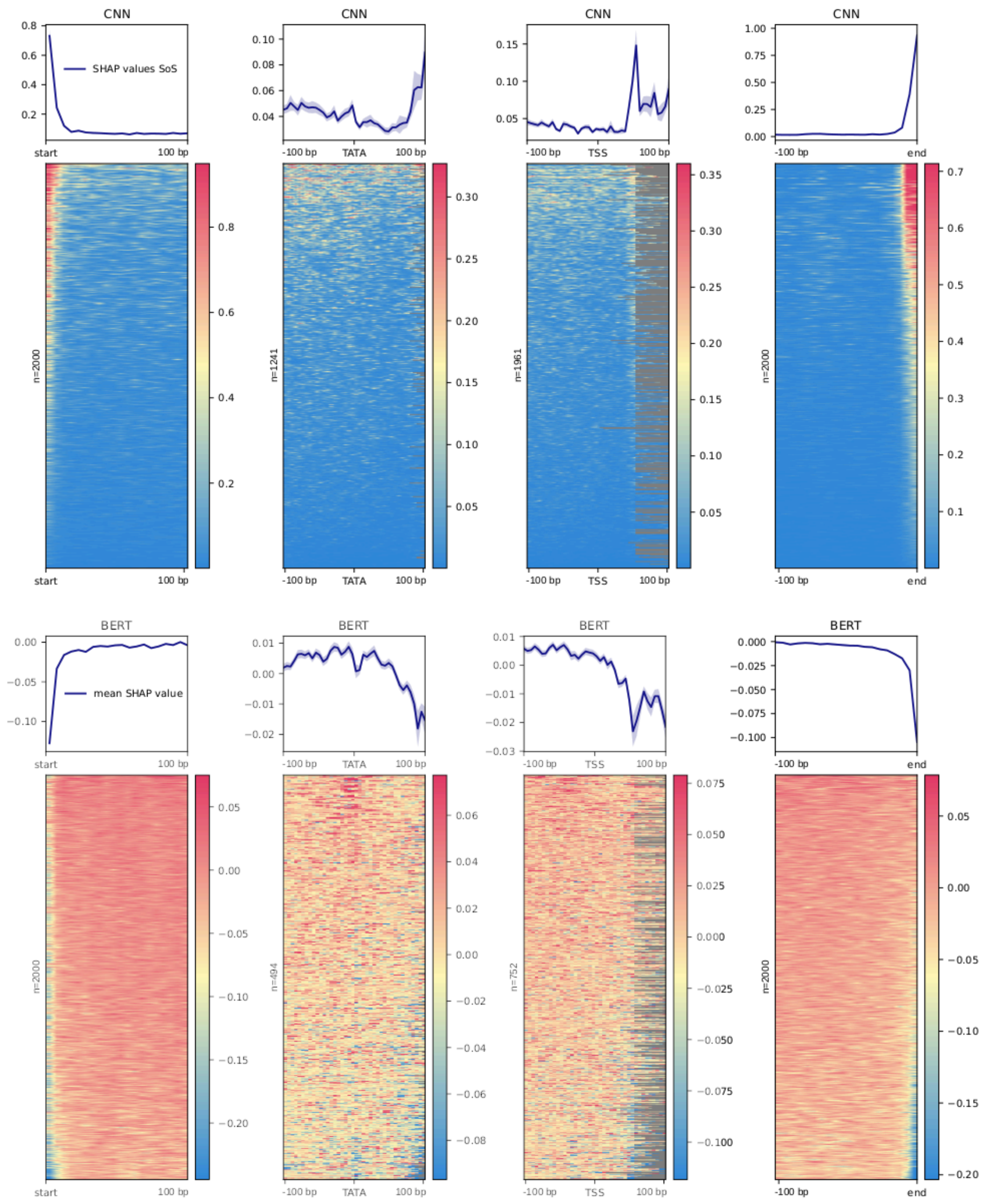


***Supplementary Figure 12 - DeepExplainer analysis of trained superfamily detection models.*** *k-mer based SHAP values were calculated along individual LTR sequences. To visualize their alignment between different sequences, sequences were aligned (from left to right) by their first base (start), predicted TATA box (TATA), predicted transcription start site (TSS), or their last base (end). Averaged SHAP values are shown as a line graph above, individual sequence values are color-coded. A - CNN model. B - DNABERT model*

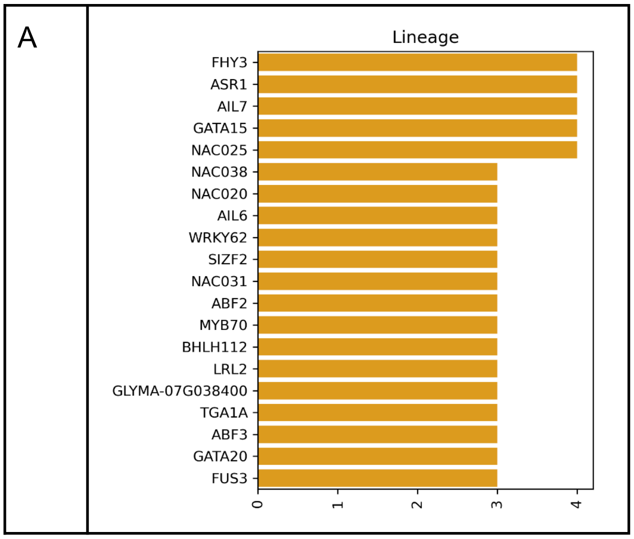


***Supplementary Figure 13 - Main results of explainability analysis carried out on trained models for family classification.*** *A - TomTom hits of first-layer CNN filters on JASPAR Core 2022 database D - B - Contribution of most significant k-mers used as input features for the DNABERT model. F - Aggregated attention heads of the DNABERT model*

Top 20 motifs by number of matches

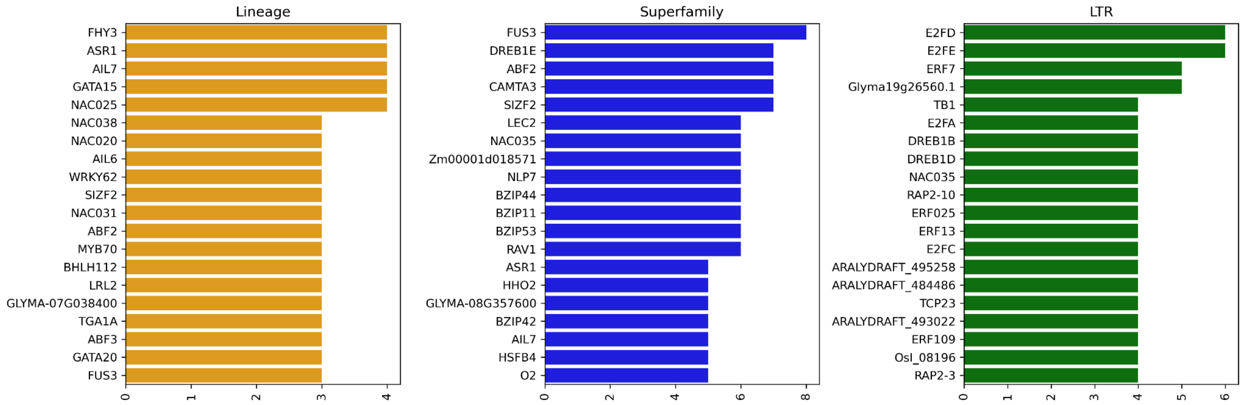


**Supplementary Figure 14 - Top TF binding sites in CNN filter analysis.** *A - Top 20 by e-value (essentially similarity of the filter to a particular binding site); B - Top 20 hits by number of filters.*

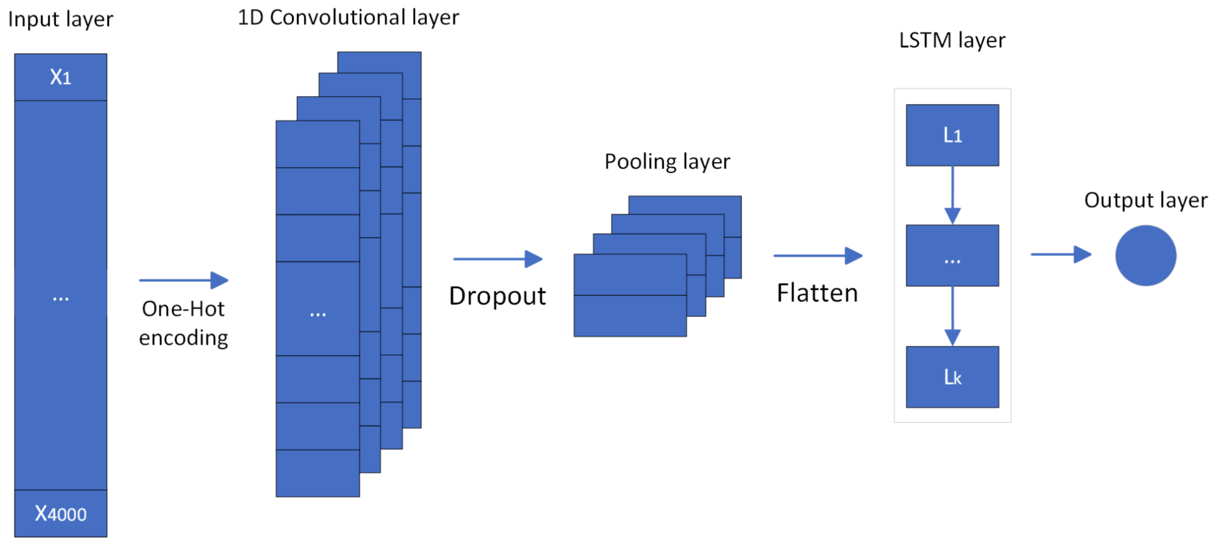


**Supplementary Figure 15 - CNN-LSTM model topology.**

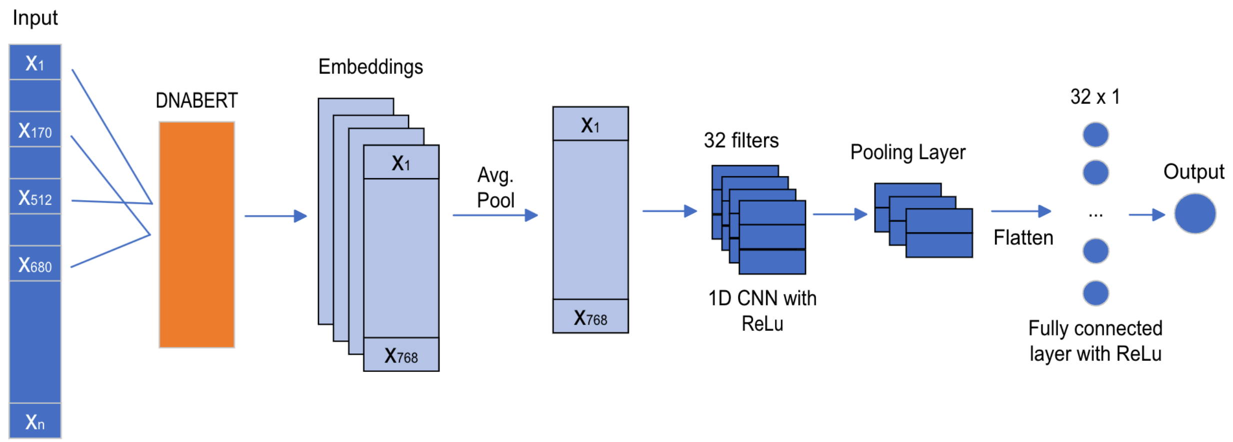


**Supplementary Figure 16 - DNABERT model topology.**

**Supplementary Tables**

| Model Type | Parameter | Tested values |
|---|---|---|
| Random Forest Classifier | N estimators | 100, 300, 600 |
| | Maximum depth | 4, 6, 8, 10, 12 |
| Multilayer Perceptron classifier | Hidden layer size | 10, 50, 100, 200 |
| | Learning rate | 0.1, 0.05, 0.02, 0.01 |
| | Alpha (Strength of L2) | 0.0001, 0.001, 0.01 |
| Gradient Boosting classifier | N estimators | 50, 100, 200, 400 |
| | Learning rate | 0.1, 0.05, 0.02, 0.01 |
| | Maximum depth | 4, 6, 8 |
| | Min samples at leaf node | 20, 50, 100, 150 |

**Supplementary Table 1 - Tested hyperparameters** during grid search for each of the three models.

| Classifier | Loss function |
|---|---|
| LTR classifier | Binary Crossentropy |
| Superfamily | Binary Crossentropy |
| Family | Categorical cross entropy |

**Supplementary Table 2 - Loss functions** used for the different classification tasks
